# Adaptive immunity to retroelements promotes barrier integrity

**DOI:** 10.1101/2024.08.09.606346

**Authors:** Alexandria C. Wells, Djalma Souza Lima-Junior, Verena M. Link, Margery Smelkinson, Siddharth R. Krishnamurthy, Liang Chi, Elisha Segrist, Claudia A. Rivera, Ana Teijeiro, Nicolas Bouladoux, Yasmine Belkaid

**Affiliations:** Metaorganism Immunity Section, Laboratory of Host Immunity and Microbiome, National Institute of Allergy and Infectious Diseases, National Institutes of Health, Bethesda, MD 20892, USA; Biological Imaging, Research Technology Branch, National Institute of Allergy and Infectious Diseases, National Institutes of Health, Bethesda, MD 20892, USA

## Abstract

Maintenance of tissue integrity is a requirement of host survival. This mandate is of prime importance at barrier sites that are constitutively exposed to the environment. Here, we show that exposure of the skin to non-inflammatory xenobiotics promotes tissue repair; more specifically, mild detergent exposure promotes the reactivation of defined retroelements leading to the induction of retroelement-specific CD8^+^ T cells. These T cell responses are Langerhans cell dependent and establish tissue residency within the skin. Upon injury, retroelement-specific CD8^+^ T cells significantly accelerate wound repair via IL-17A. Collectively, this work demonstrates that tonic environmental exposures and associated adaptive responses to retroelements can be coopted to preemptively set the tissue for maximal resilience to injury.

## Main text

Barrier tissues, of which the skin is the largest and outermost, serve as a primary interface with the environment. As such, this tissue is constitutively exposed to environmental stressors ranging from innocuous exposures to severe damages caused by infection, inflammation, or injury (*1, 2*). In regards to the latter, protective responses must be quickly and efficiently initiated to maintain this barrier site. Whether mild environmental stressors can preventively prepare tissues for maximal resilience remains unclear.

Preservation of host integrity is one of the dominant forces driving evolution; thus, we speculate that all components of the system, both host-derived and acquired, may converge toward this fundamental function. Further, sophistication of the host throughout evolution implies that multiple systems may have been superimposed to repair highly complex tissues. This concept is supported by emerging work from us and others uncovering a role for both adaptive immunity and the microbiota in the promotion of tissue repair (*1, 3, 4*). Notably, within the skin, microbiota-reactive T cells have been implicated in mediating wound repair (*5–9*). How additional elements acquired through evolution further synergize to preserve barrier integrity remains an outstanding question.

An additional layer of genetic material that has progressively integrated into the host genome are endogenous retroelements (EREs): genomic parasites that rely on reverse transcription and integration of their genomes into host DNA, and comprise 43% of the human genome and 37% of the mouse genome (*10, 11*). These elements play essential roles in diverse processes, including placental morphogenesis, mammalian development, evolution of the V(D)J gene segments of B and T cell receptors, and orchestrating the afore-mentioned relationship with the microbiota (*8, 12, 13*). Moreover, throughout their ‘lifecycle’ EREs generate many nucleic acid intermediates and proteins that can engage both the innate and adaptive immune systems, respectively (*8, 14–16*). As such, EREs may represent one of the dominant sources of “self-antigen” in the host. Indeed, EREs have been shown to be a source of immunogenic antigens under defined pathogenic settings such as cancer, autoimmune diseases, or infection (*14, 15*). Whether EREs can be directly and specifically recognized by the adaptive immune system under non-pathogenic and/ or homeostatic conditions, and the significance of these responses for host physiology remains unclear. Here, we tested the possibility that non-pathogenic environmental exposures may promote ERE-specific immune responses and uncovered a role for adaptive immunity to EREs in the promotion of tissue resilience in the face of barrier damage.

### Mild detergent exposure promotes CD8^+^ T cell responses in the skin

To test tissue responses to mild environmental stressors, we developed a model of detergent exposure where we topically applied mild detergents containing brain heart infusion broth emulsified in either 1% tween-80 (BHI-tween) or sodium dodecyl sulfide (BHI-SDS) (**Fig. 1A**). One such treatment did not cause tissue inflammation, as there was neither appreciable thickening of the skin, nor accumulation of neutrophils, monocytes and basophils (**fig. S1A, B**). On the other hand, both detergents induced a significant increase in CD8^+^ T cell frequency and numbers (**Fig. 1A, fig. S1C**), with minimal accumulation of CD4^+^, TCRγδ T cells, and mucosal-associated invariant T (MAIT) cells (**Fig. 1B, fig. S1D-F**). Because a more significant number of CD8^+^ T cells accumulated in response to BHI-tween, we subsequently focused on this specific regiment.

**Fig. 1.**
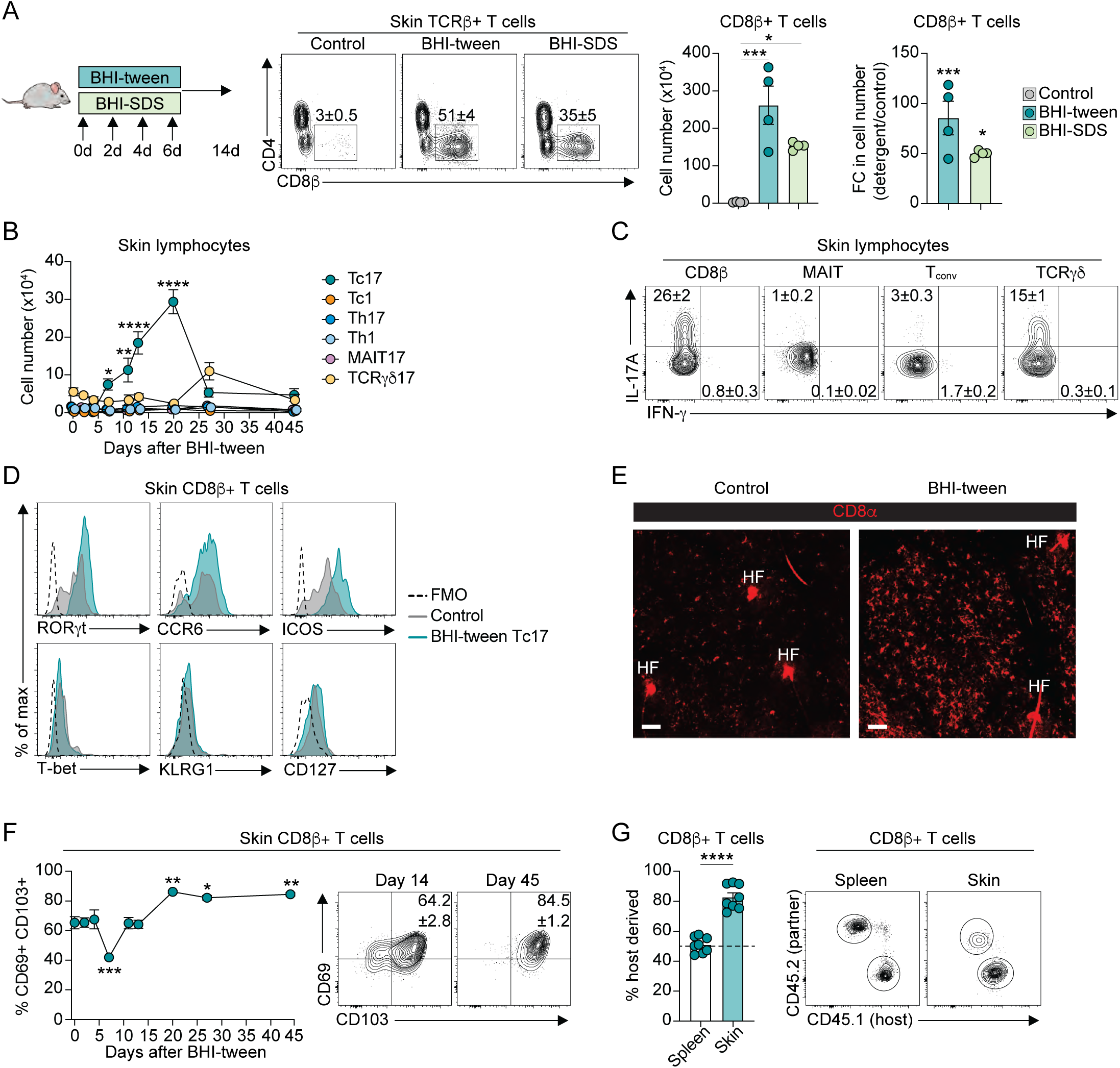
Environmental exposures recruit CD8^+^ T cells to the skin. **(A)** Left, experimental design of detergent exposure. Middle, representative FACS plots showing frequencies of CD8β^+^ T cells in ear pinnae skin of unexposed (control), or detergent exposed (BHI-tween, BHI-SDS) animals. Right, quantification of CD8β^+^ T cells in (left) absolute numbers or (right) as fold change in absolute numbers in detergent exposed mice compared to control animals. **(B)** Absolute numbers of IL-17A- and IFN-γ-producing lymphocytes in ear pinnae skin at the indicated time after BHI-tween exposure. Lymphocytes shown are CD8β^+^ T cells, Mucosal-associated Invariant T cells (MAIT), CD4^+^FoxP3^-^ helper T cells (Th), TCRγδ-mid T cells. **(C)** Representative FACS plots showing the frequencies of IL-17A- and IFN-γ-producing lymphocytes in ear pinnae skin 14 days after BHI-tween exposure. **(D)** Representative histograms showing expression of RORγt, CCR6, ICOS, T-bet, KLRG1, and CD127 by Tc17 cells in ear pinnae skin of mice 14 days after BHI-tween exposure, or total CD8β^+^ T cells in ear pinnae skin of control animals. Dashed gray line represents fluorescence minus one (FMO) controls. **(E)** Representative whole mount confocal microscopy images of CD8β^+^ T cells in the ear pinnae skin of control animals or mice 14 days after BHI-tween application. Scale bars are 50 μm. HF, hair follicle **(F)** (Left) frequencies or (right) representative FACS plots showing frequencies of CD69^+^ CD103^+^ CD8β^+^ T cells in ear pinnae skin at the indicated time after BHI-tween exposure. **(G)** (Left) frequencies or (right) representative FACS plots of host-derived CD8β^+^ T cells in the spleen or flank skin of parabiotic mice, as indicated by either CD45.1^+^ or CD45.2^+^ cells within the CD8β^+^ T cell compartment. (A-G) Data are representative of at least two independent experiments. (A, G) Each dot represents an individual mouse. (B, F) Each dot represents the mean of five individual mice. Numbers in representative flow plots indicate mean ± SEM. * p < 0.05; ** p < 0.01; *** p < 0.001; **** p < 0.0001; ns, not significant (One-way ANOVA with Dunnett’s multiple comparison test (compared to control or day 0) for A, B, F; two-tailed unpaired Student’s t test for G).

CD8^+^ T cells induced by mild detergent predominantly produced IL-17A (Tc17), with few IFN-γ producers (Tc1) (**Fig. 1B, C**). Accordingly, detergent-induced Tc17 cells highly expressed canonical type 17 markers including RORγt, CCR6, and ICOS, and did not express type 1 markers (T-bet, KLRG1, and CD127) (**Fig. 1D**). Tc17 responses to mild detergent were also detected in the regional lymph node (rLN), with only a small expansion detectable in the spleen (**fig. S1G**).

In a comparable manner to CD8^+^ T cell responses to viral infections (*17–19*), detergent-induced Tc17 cells were enriched within the epidermis, where they expressed a higher level of IL-17A and RORγt than in the dermis (**fig. S1H, I**). Confocal imaging revealed that detergent-induced CD8^+^ T cells were evenly distributed throughout the tissue (**Fig. 1E**).

Previous literature uncovered that CD8^+^ T cells recruited to the epidermis can develop as tissue resident T cells (*20, 21*). Kinetic assessment demonstrated that following exposure to detergent, Tc17 cells began to accumulate at day 7, peaked at day 21, and following contraction at day 28, were still detected up to 45 days post-detergent application (**Fig. 1B**). Further, Tc17 cells expressing tissue residency markers (CD103 and CD69) were enriched following the peak of the response (**Fig. 1F**). To test the possibility that these cells could develop as *bona fide* tissue resident cells, we generated congenically-mismatched parabiotic pairs with mice that had been exposed to mild detergent 30 days prior (**fig. S1J**). By 30 days post-joining, CD8^+^ T cells within the skin were primarily host-derived, as compared to splenic CD8^+^ T cells which were 50% host-derived (**Fig. 1G**). Consistent with a recent report demonstrating reduced durability of skin tissue-resident Tc17 cells (T_RM_17) compared to tissue-resident Tc1 cells (T_RM_1) (*22*), detergent-induced Tc17 cells were not fully stable, with 83% of Tc17 being host-derived (**Fig. 1G**). Thus, detergent-induced CD8^+^ T cells can develop as tissue-resident Tc17 cells within the skin. As such, exposure to mild environmental stressors influences the tissue immune landscape, in particular via the accumulation of tissue-resident Tc17 cells.

### Endogenous retroelements are reactivated in Langerhans cells following detergent exposure

We and others have shown that defined members of the skin microbiota can promote Tc17 responses (*5, 23, 24*); however, detergent-induced CD8^+^ T cell responses were still detected in mice devoid of live microbes (GF) (**Fig. 2A**). This observation raised the intriguing possibility that these xenobiotic-induced responses were self-reactive.

**Fig. 2.**
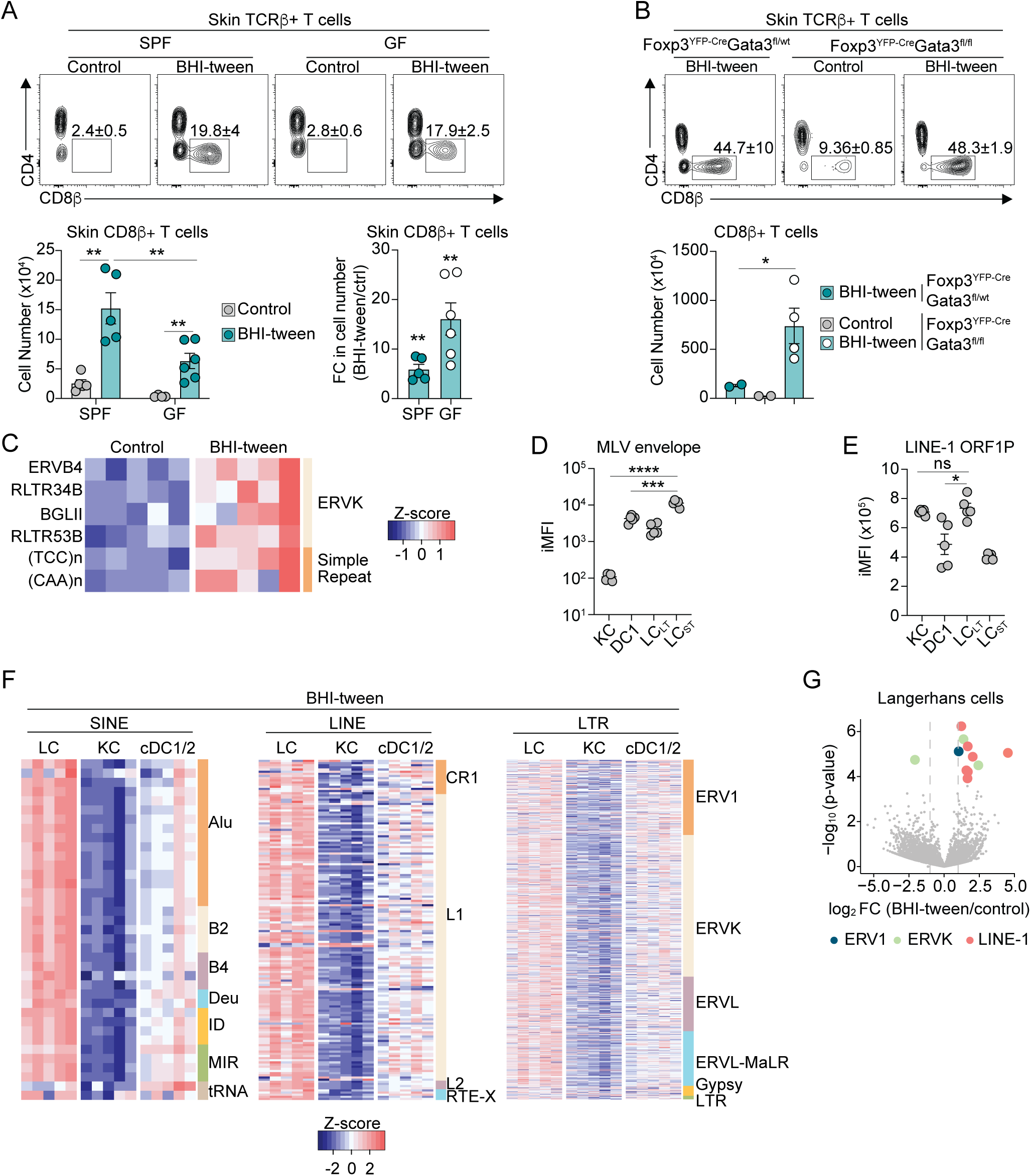
Detergent exposure reactivates expression of endogenous retroelements. **(A)** (Top) representative FACS plots showing frequencies, or (bottom left) absolute numbers of CD8β^+^ T cells in ear pinnae skin of specific pathogen-free (SPF) or germ-free (GF) mice 14 days after BHI-tween exposure or of control animals. Bottom right, fold change in absolute numbers of CD8β^+^ T cells in ear pinnae skin of SPF or GF mice 14 days after detergent exposure compared to control SPF or GF mice, respectively. **(B)** (Top) representative FACS plots showing frequencies or (bottom) absolute numbers of CD8β^+^ T cells in ear pinnae skin of control or detergent exposed Foxp3^YFP-Cre^Gata3^fl/wt^ (wild type) or Foxp3^YFP-Cre^Gata3^fl/fl^ animals. **(C)** Heatmap displaying fold change of significantly differentially expressed retroelement elements (FC > 2, FDR < 0.05) by RNA-sequencing of whole homogenized ear pinnae skin from control mice or animals exposed to BHI-tween daily for 7 days. Retroelement elements are listed on the left, and retroelement classes on the right. **(D-E)** Protein abundance displayed as Integrated MFI (iMFI) of the (D) MLV envelope or (E) LINE-1 ORF1p by flow cytometry in keratinocytes, conventional dendritic cells type 1 (cDC1), long-term Langerhans cells (LC_LT_, EpCAM^hi^), or short-term Langerhans cells (LC_ST_, EpCAM^lo^) in control animals. **(F)** Heatmap displaying row z-score of retroelement elements from RNA-sequencing of detergent-exposed Langerhans cells (LC), keratinocytes (KC; Sca-1^+^, CD49f^+^), and conventional dendritic cells (cDC1/2) sort-purified from ear pinnae skin exposed daily to BHI-tween for 7 days. (Left) Short-Interspersed Nuclear Elements (SINE), (middle) Long-Interspersed Nuclear Elements (LINE), (right) Long Terminal Repeats (LTR). Retroelement families are listed on the right. Deu, deuterostome; ID, identifier; MIR, mammalian-wide interspersed repeat; RTE, retrotransposable element clade; ERV, endogenous retrovirus; MaLR, mammalian apparent LTR retrotransposon. **(G)** Volcano plot displaying differentially expressed retroelement loci identified by RNA-sequencing in bulk Langerhans cells sort-purified from ear pinnae skin of mice exposed to BHI-tween daily for 7 days as compared to control mice. (A, B, D, E) Data are representative of at least two independent experiments. Each dot represents an individual mouse. Numbers in representative flow plots indicate mean ± SEM. * p < 0.05; ** p < 0.01; *** p < 0.001 (two-way ANOVA with Tukey’s multiple comparisons test for A; two-tailed unpaired Student’s t test for B, D, E).

In line with this possibility, reducing local regulatory T cell (T_reg_) activity by specifically deleting GATA3 from Foxp3-expressing cells (GATA3^+^ T_regs_ are the majority of cutaneous T_regs_), led to an increase in the xenobiotic-induced CD8^+^ T cell response (**Fig. 2B**) (*6, 25–27*). This included Tc17, as well as an accumulation of Tc1 not typically observed in the response to mild detergent (**Fig. 1B, fig. S2A**). Further, in mice deficient in GATA3^+^ T_regs_, detergent exposure was associated with local inflammation, as evidenced by an increase in neutrophils and monocytes as compared to control mice (**fig. S2B**). This supported the idea that the pathogenic potential of these responses was under constitutive control of local regulatory T cells.

Endogenous retroelements (EREs) constitute 37% of the mouse genome (and 43% of the human genome) and thus represent an enormous potential source of self-antigen. While responses to EREs have been observed in defined pathogenic states (*14*), whether EREs can be reactivated and recognized by the adaptive immune system during homeostasis or following mild xenobiotic exposure remains unclear. Analysis of ERE expression within the skin following detergent application revealed a discreet, yet discernable, alteration in ERE expression (**Fig. 2C**). Of note, ERE elements that were expressed following detergent application were distinct from those induced by *S. epidermidis* (**fig. S2C**) (*8*), supporting the idea that defined environmental exposures promote distinct ERE expression patterns.

To assess whether ERE activity may be enriched within defined antigen presenting cell subsets, we first quantified the expression of the envelope protein of Murine Leukemia Virus (MLV) and ORF1p of Long-Interspersed Nuclear Elements-1 (LINE-1) in distinct cell subsets at baseline (**Fig. 2D, E, fig. S2D**). In particular, two subsets of Langerhans cells (LCs) had the highest expression of LINE-1 ORF1p and MLV envelope proteins, respectively, as compared to keratinocytes (KCs) and conventional DC1s (cDC1s) (**Fig. 2D, E**). These subsets either represent embryonic-derived LCs (long-term LC, LC_LT_) or the bone-marrow derived LC precursors which replace LC_LT_ that have emigrated out of the tissue (short-term LC, LC_ST_) (*28–30*). Thus, at baseline, Langerhans cells have elevated ERE-derived protein expression compared to other cell subsets.

Bulk RNA-seq analysis of LCs, cDCs (pooled cDC1 and cDC2), and keratinocytes purified from naïve or detergent exposed mice was used to gain a deeper perspective of which retroelement species were reactivated in this setting. In support of our flow cytometry data, LCs from naïve mice expressed significantly higher levels of most elements associated with a given ERE family, as compared to cDCs and keratinocytes, further suggesting there is constitutively high ERE expression in this cell subset (**fig. S2E**).

Elevated expression of most ERE families in LCs, as compared to keratinocytes and cDCs, was also observed following detergent exposure (**Fig. 2F**). At the individual loci level, LINE-1 elements were the most abundant significantly upregulated EREs in detergent-exposed LCs as compared to unexposed LCs (**Fig. 2G**). Albeit at a lower level than LCs, keratinocytes and cDCs also increased ERE expression post-detergent exposure (**fig. S2F**). Interestingly, the majority of loci upregulated by detergent exposure in keratinocytes were Endogenous Retrovirus K (ERVK), while cDCs equally upregulated ERV1 and LINE-1 loci, further highlighting the distinct pattern of expression between cell subsets.

Thus, Langerhans cells, which represent the most abundant antigen-presenting cell (APC) in the skin, have elevated basal expression of EREs compared to other cutaneous APCs, and further upregulate these elements in response to mild detergent. These results propose that Langerhans cell-intrinsic EREs may represent a source of antigen for CD8^+^ T cell responses in the context of xenobiotic exposure.

### Langerhans cells are required for CD8^+^ T cell responses to detergent exposure

We next explored the potential role of LCs in driving the CD8^+^ T cell response to mild detergent. In agreement with the involvement of skin APCs, T cell responses were abrogated in mice in which DC migration, including LCs, to the regional lymph node is impaired (CCR7-deficient mice) (**Fig. 3A** and **fig. S3A**). We also found that the number of LCs was significantly increased within the regional lymph node (rLN), reaching peak accumulation 8 days post-detergent exposure, immediately preceding the peak of CD8^+^ T cell accumulation in the skin (**Fig. 3B** and **fig. S3B**). On the other hand, cDC1s did not appreciably accumulate in the rLN until 12 days after the initial detergent application, a time past the initiation of the CD8^+^ T cell response (**Fig. 3B, fig. S3B and Fig. 1B**). In line with the non-inflammatory nature of these responses, peak APC accumulation in the rLN was delayed compared to what has been described in the context of infection (*31*).

**Fig. 3.**
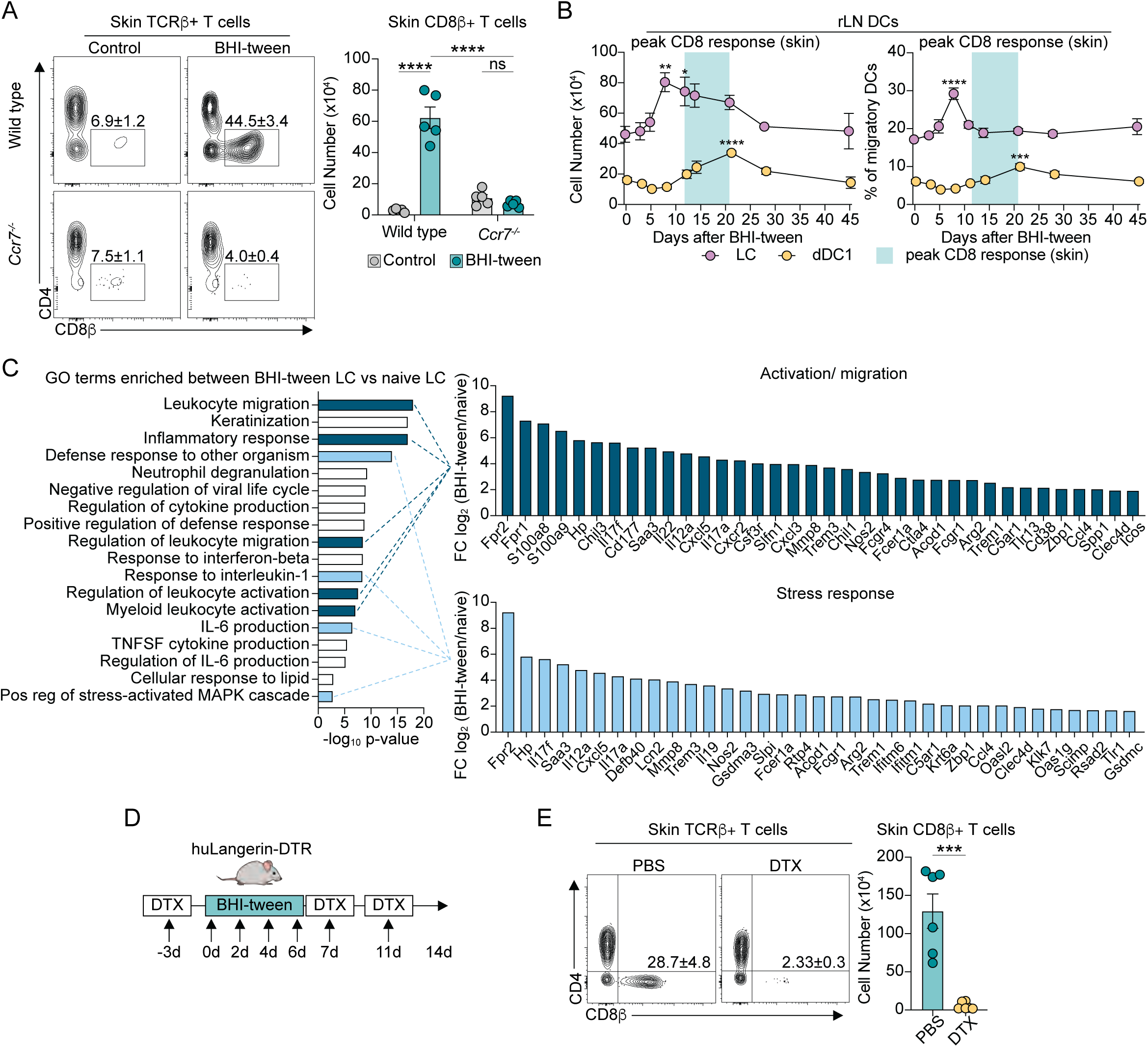
Langerhans cells are required for CD8^+^ T cell responses to mild detergent. **(A)** (Left) representative FACS plots showing the frequencies or (right) absolute numbers of CD8β^+^ T cells in ear pinnae skin of wild type or *Ccr7^-/-^* animals either as a control or exposed to BHI-tween 14 days prior. **(B)** (Left) absolute numbers of migratory dDC1 or Langerhans cells, or (right) frequencies of dDC1 and Langerhans cells among all migratory DCs in the ear pinnae-regional lymph node at the indicated time after BHI-tween exposure. Peak accumulation of CD8β^+^ T cells in the skin is highlighted. **(C)** (Left) top 18 GO terms enriched in Langerhans cells from mice exposed to BHI-tween daily for 7 days. (Right) top 35 genes enriched in detergent-exposed Langerhans cells related to Langerhans cell (top) activation/ migration or (bottom) immune stress response. **(D)** Experimental design of diphtheria toxin treatment to deplete Langerhans cells throughout the response to BHI-tween exposure. DTX, diphtheria toxin. **(E)** (Left) representative FACS plots showing the frequencies or (right) absolute numbers of CD8β^+^ T cells in ear pinnae skin 14 days after BHI-tween treatment of huLangerin-DTR mice treated either with PBS (Langerhans cells sufficient) or diphtheria toxin (DTX, Langerhans cells depleted). (A, B, D) Data are representative of at least two independent experiments. Each dot represents an individual mouse. Numbers in representative flow plots indicate mean ± SEM. *** p < 0.001; **** p < 0.0001; ns, not significant (two-way ANOVA with Tukey’s multiple comparisons test for A; one-way ANOVA with Dunnett’s multiple comparison test (compared to day 0) for B; two-tailed unpaired Student’s t test for D).

We next assessed the transcriptional response of LCs to detergent exposure. Functional enrichment analysis confirmed that Langerhans cells were activated upon detergent exposure, as demonstrated by elevated expression of genes associated with the GO terms “leukocyte migration” and “inflammatory response”, including *Fpr2*, *S100a8*, *Chil3*, *Cxcl5*, *Cxcr2*, *Mmp8* and *Nos2* (**Fig. 3C**). Other transcripts induced in detergent-exposed LCs were related to immune cell activation in response to stress. One of the most upregulated genes was serum amyloid A (*Saa3*) (**Fig. 3C**), a gene whose expression is associated with the acute phase response to various physiological stressors, such as infection or injury (*32, 33*). In addition, genes associated with the response to reactive oxygen species (*Nos2, Il19*), IL-1β and IL-6 (*Acod1, Il19, Scimp*), type I interferons (*Ifitm6, Ifitm1, Oasl2, Oas1g*), and DAMP/PAMP recognition (*Tlr1, C5ar1*) were upregulated, emphasizing the profound impact detergent exposure has on the activation status of LCs in the skin (**Fig. 3C**).

To specifically test the role of LCs in xenobiotic-induced CD8^+^ T cell responses, we employed an approach to selectively deplete LCs while sparing Langerin^+^ dermal DC1 **(Fig. 3D** and **fig. S3C**) (*34, 35*). Detergent-exposed huLangerin-DTR mice were treated with diphtheria toxin (DTX) prior to detergent exposure (**Fig. 3D**). Strikingly, detergent-induced CD8^+^ T cell accumulation was entirely abrogated in the absence of LCs, revealing an essential role for LCs in driving these responses (**Fig. 3E**). Taken together, our results suggest LCs transcriptionally respond to detergent exposure and are required for CD8^+^ T cell responses.

### Detergent-exposed CD8^+^ T cells recognize LINE-1-derived antigens

We next explored the nature of detergent-induced CD8^+^ T cells. CD8^+^ T cell responses to mild detergent were abrogated in β-2m-related MHC-I-deficient animals (**fig. S4A**), eliminating the possibility that these cells were non-classical intraepithelial CD8^+^ T cells (*36*). Additionally, CD8^+^ T cell responses to detergent were not impacted in the absence of H2-M3 (**fig. S4B**), the MHC-Ib molecule that presents *N*-formyl methionine-led peptides, further suggesting that these cells were not microbiota-reactive (*5*).

We next tested the hypothesis that LCs may be the relevant source of antigen for CD8^+^ T cells by leveraging an *in silico* approach to identify LC-intrinsic ERE-derived peptides (**Fig. 4A**). Specifically, using the previously generated ERE RNA-seq dataset from sort-purified naïve and detergent-exposed LCs, cDCs, and keratinocytes, we identified 10 loci that were expressed at a minimum of 4 transcript per million (TPM) and upregulated in response to mild detergent exposure in LCs (fold change > 2, false discovery rate < 0.05) (**Fig. 4B, table S1**). Confirming the previous analysis, 80% of these loci were uniquely expressed in LCs, except for Chr.5 2.37- and Chr.17 3.23+ which were also expressed in keratinocytes and cDCs. Furthermore, these 10 loci were cross-referenced and filtered for expression in medullary thymic epithelial cells (mTECs), which mediate negative selection of autoreactive CD8^+^ T cells (*10, 15, 37*). Of note, only Chr.5 2.37- and Chr. 17 3.23+ were expressed in mTECs (**Fig. 4B**) and were thus excluded from downstream peptide screening.

**Fig. 4.**
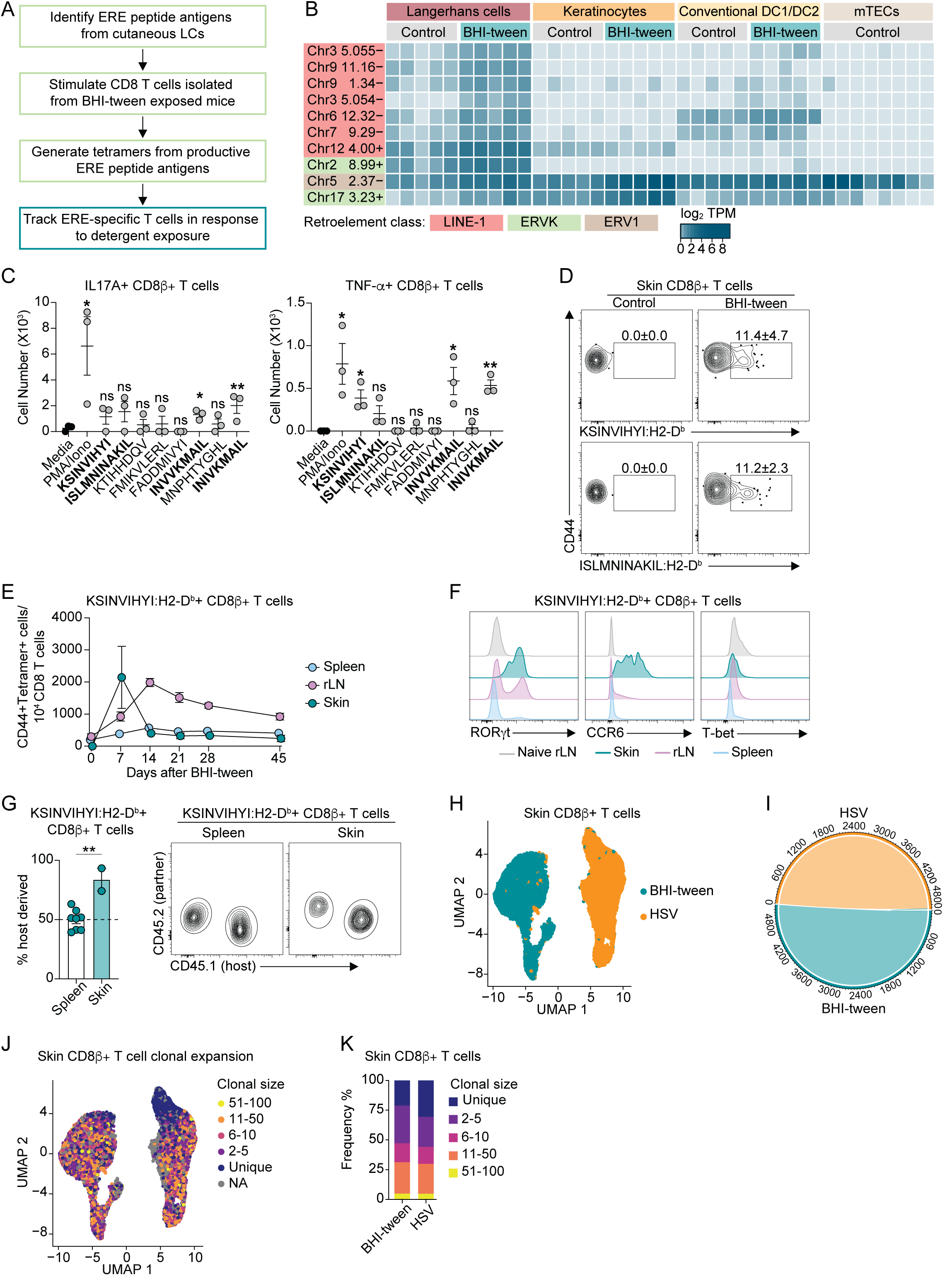
LINE-1-specific CD8+ T cells clonally expand in response to detergent exposure. **(A)** Pipeline used to generate tetramers to identify ERE-specific CD8β^+^ T cells. **(B)** Heatmap displaying expression of retroelement loci from RNA-sequencing of bulk Langerhans cells, keratinocytes (Sca-1^+^, CD49f^+^), and conventional dendritic cells (cDC1/2) sort-purified from ear pinnae skin exposed daily to BHI-tween for 7 days. Medullary Thymic Epithelial Cells (mTECs) were sort-purified from unexposed wild type mice, data was extracted from (Cowan 2019 NC). LINE-1: Long-Interspersed Nuclear Element-1, ERVK: Endogenous Retrovirus-K, ERV-1: Endogenous Retrovirus-1. **(C)** Absolute numbers of CD8β^+^ T cells producing either (left) IL-17A or (right) TNF-α in response to *ex vivo* stimulation with dendritic cells from unexposed control mice pulsed with ERE-derived peptides. Peptides in bold were selected to generate tetramers. **(D)** FACS plots showing the frequencies of KSINVIHYI:H2-Db^+^ or ISLMNINAKIL:H2-D^b+^ CD8β^+^ T cells in the ear pinnae skin of control animals or mice exposed to BHI-tween 14 days prior. Plots are concatenated from three mice. **(E)** Number of CD44^+^ KSINVIHYI:H2-D^b+^ CD8β^+^ T cells relative to total CD8β^+^ T cells in ear pinnae skin, ear pinnae-regional lymph node (rLN), and spleen at the indicated time after BHI-tween exposure. **(F)** Representative histograms showing expression of RORγt, CCR6, and T-bet by KSINVIHYI:H2-D^b+^ CD8β^+^ T in ear pinnae skin, ear pinnae-regional lymph node, and spleen of mice 14 days after BHI-tween exposure, or the rLN of control animals. **(G)** (Left) frequencies or (right) representative FACS plots showing frequencies of host-or donor-derived KSINVIHYI:H2-D^b+^ CD8β^+^ T cells, as indicated by either CD45.1^+^ or CD45.2^+^, in the spleen or ear pinnae skin of parabiotic mice. **(H)** UMAP projection plot showing CD8β^+^ T cells sort-purified from ear pinnae skin exposed to BHI-tween 14 days prior or 7 days post-HSV-1 infection by scRNA-seq. **(I)** Circos plot showing clonal sharing between treatment groups. **(J)** UMAP projection plot or **(K)** bar plot showing the relative abundance of CD8β^+^ T cell clonotype expansion (determined by TCR amino acid sequence via scRNA-seq). (C-G) Data are representative of at least two independent experiments. Numbers in representative flow plots indicate mean ± SEM. (C) Each dot represents a technical replicate. (E) Each dot represents the average of 5 mice. (G) Spleen, each dot represents an individual mouse. Skin, each dot represents 4 concatenated mice. * p < 0.05; ** p < 0.01; *** p < 0.001; **** p < 0.0001; ns, not significant (two-tailed unpaired Student’s t test for C, G).

Four peptides (KSINVIHYI, ISLMNINAKIL, INVKKMAIL, INIVKMAIL), all derived from LINE-1 loci (**table S1**), triggered either IL-17A or TNF-α production (**Fig. 4C** and **fig. S4C**). To verify *in vivo* reactivity and track these responses, we generated tetramers, and validated their specificity by dual fluorophore tetramer staining (**fig. S4D**). As all SPF animals are naïve to the bacterial pathogen *Yersinia pseudotuberculosis*, the inability of the YopE antigen tetramer (SVIGFIQRM:H2-K^b^) to label detergent-elicited CD8^+^ T cells was used to further confirm the specificity of the LINE-1 tetramers (**fig. S4D**).

At the peak of the CD8^+^ T cell response to detergent, KSINVIHYI and ISLMNINAKIL positively labeled 4% of CD8^+^ T cells within the skin, indicating that LINE-1-specific CD8^+^ T cells are indeed a component of this response (**Fig. 4D** and **fig. S4E**). LINE-1-specific CD8^+^ T cell responses were highly enriched at the site of exposure, and expansion of LINE-1-specific CD8^+^ T cells occurred to a higher magnitude in the rLN than in the spleen (**Fig. 4E** and **fig. S4E**). Further, and in agreement with the polyclonal response, LINE-1-specific CD8^+^ T cells displayed a Tc17 phenotype (RORγt^+^, CCR6^+^, T-bet^-^) in the skin and rLN, and to a lesser extent in the spleen (**Fig. 4F**).

In the skin, KSINVIHYI:H2-D^b+^ CD8^+^ T cells could be detected as far as 45 days following the initial detergent exposure (**Fig. 4E** and **fig. S4E**). In a similar manner to the polyclonal Tc17 response (**Fig. 1F**) parabiosis experiments revealed that LINE-1-specific CD8^+^ T cells also formed a *bona fide* tissue resident population (**Fig. 4G**), with 80% of cells residing stably in the skin, a property reflective of their type 17 signature (**fig. S4F**).

As LINE-1-specific CD8^+^ T cells represented a substantial proportion of the mild detergent response, we next assessed the diversity of the detergent-responsive CD8^+^ T cell TCR repertoire. To this end, we performed single cell RNA-sequencing analysis on detergent-induced CD8^+^ T cells and compared them to the well-characterized CD8^+^ T cells that undergo clonal expansion in response to Herpes Simplex Virus-1 (HSV-1) infection (*21*). As expected, CD8^+^ T cells clustered based on their treatment in UMAP analysis (**Fig. 4H**). Each replicate mouse contributed a similar frequency of unique clonotypes to the data and were thus representative of each treatment (**fig. S4G**). Additionally, only one clone overlapped between conditions (**Fig. 4I**), supporting the idea that there were no public TCRs shared among treatments.

Interestingly, both treatment groups generated responses with similar frequencies of hyper and large expanded clonotypes (4.8% and 26.4% in BHI-tween, and 4.7% and 25.2% in HSV-1, respectively) (**Fig. 4J, K**), despite high variability in the putative peptide antigens presented in these settings. This supported the idea that while LINE-1-specific CD8^+^ T cells were a notable fraction of the total CD8^+^ T cell pool, the xenobiotic-induced CD8^+^ T cell response is in fact highly polyclonal. Overall, these data support the idea that detergent exposure induce CD8^+^ T cell responses to ERE antigens.

### Xenobiotic-induced CD8^+^ T cells are enriched for wound repair signature

To further characterize detergent-elicited CD8^+^ T cells, we analyzed their gene expression by single cell RNA-sequencing. As expected, detergent-elicited CD8^+^ T cells were transcriptionally distinct from their anti-viral counterparts (**Fig. 4H**) and in contrast with HSV-1-induced CD8^+^ T cells, were dominated by a Tc17 signature (**fig. S5A-C**). Based on previously established markers (*38, 39*), the detergent-induced CD8^+^ T cell response included Tc17 cells at various stages of activation (**Fig. 5A**). To explore the potential function of detergent-reactive CD8^+^ T cells, gene expression of detergent-elicited Tc17 and HSV-1-induced Tc1 cells was assessed by bulk RNA-seq. Accordingly, CD8^+^ T cells responding to mild detergent highly expressed genes associated with a type 17 signature, including *Ccr6*, *Il17a*, *Il17f*, and *Rorc* (**Fig. 5B**). Notably, genes coding for other nuclear receptor transcription factors (*Rora*, *Nr4a1*), and genes involved in epithelial and vascular growth (*Areg*, *Furin*, *Itgav*, *Angptl2*) were also strongly upregulated in Tc17 cells (**Fig. 5C**).

**Fig. 5.**
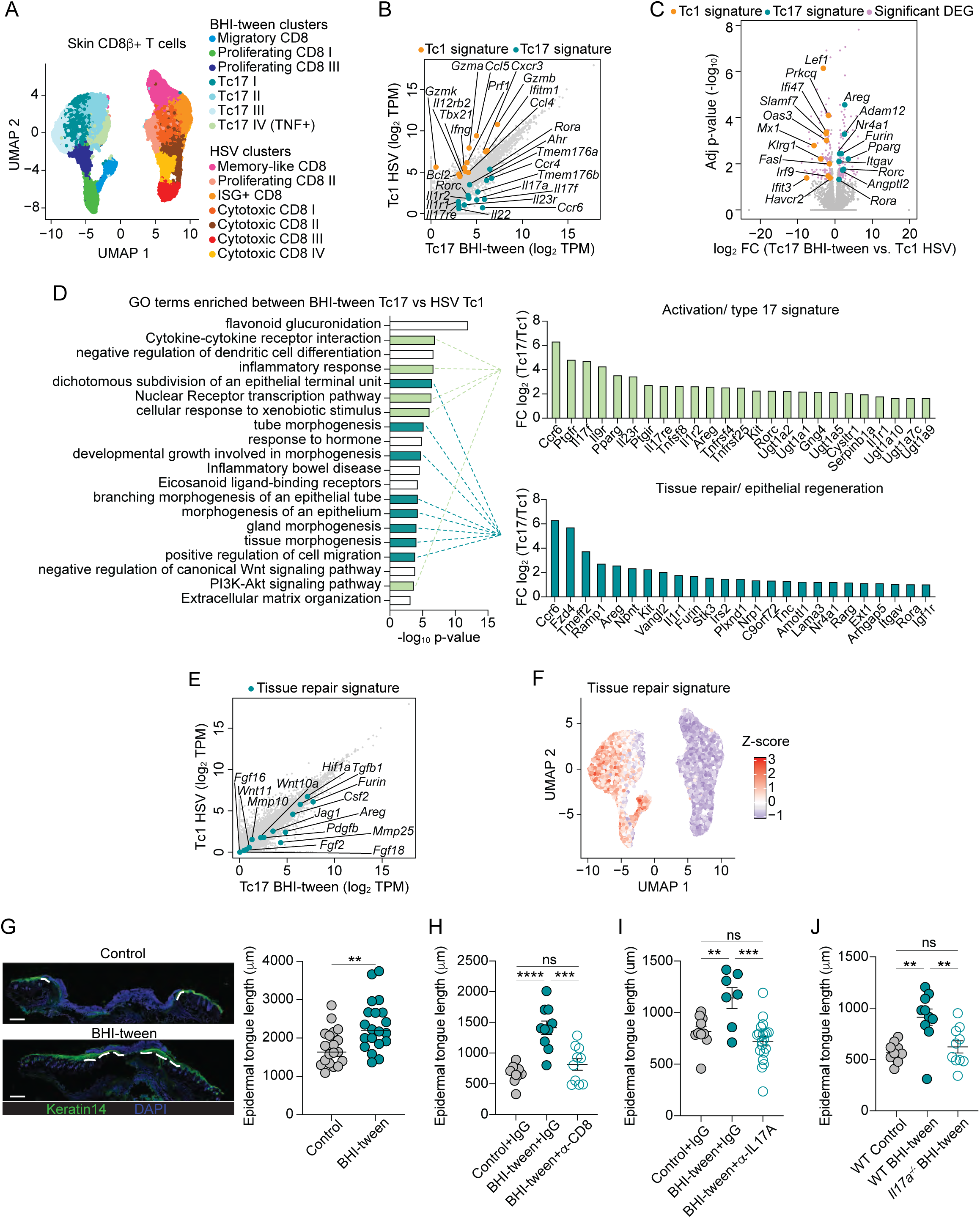
Detergent-induced CD8^+^ T cells are enriched for a repair signature and accelerate wound repair. **(A)** UMAP projection plot showing ear pinnae skin CD8β^+^ T cells sort-purified from animals 14 days post-BHI-tween exposure or 7 days post-HSV-1 infection by scRNA-seq. (**B-E**) CCR6^-^ CD8β^+^ T cells (Tc1) or CCR6^+^ CD8β^+^ T cells (Tc17) were sort-purified from ear pinnae skin of HSV-1-infected or BHI-tween-exposed animals, respectively, for bulk RNA-sequencing analysis. (B) Scatterplot showing genes related to type 1 and type 17 signatures in Tc1 and Tc17 cells. (C) Volcano plot showing differentially expressed genes comparing Tc1 and Tc17 cells. (D) (Right) top 20 GO terms enriched in Tc17 compared to Tc1 cells, or (left) top 25 genes enriched in detergent-exposed Tc17 cells related to (top) activation/ type 17 signature and (bottom) tissue repair/ epithelial regeneration. (E) Scatterplot showing genes related to tissue repair in Tc1 and Tc17 cells. **(F)** UMAP projection plot showing the wound repair signature by scRNA-seq in CD8β^+^ T cells sort-purified from ear pinnae skin of animals 14 days post-BHI-tween exposure or 7 days post-HSV-1 infection. **(G-J)** Mice were untreated or exposed to BHI-tween 12 days prior to back skin punch biopsy, and epithelial tongue length measured 5 days after wounding. (G) (Left) representative immunofluorescence images or (right) quantification of epidermal tongue length. Tissue sections are stained for basal keratinocytes (keratin 14, green) and DAPI (blue). White dashed lines represent the epidermal tongue length during re-epithelialization of the wound. Scale bars are 500 μm. (H) Quantification of the epidermal tongue length in control mice or BHI-tween exposed mice treated with either an isotype control antibody (IgG) or an anti-CD8α depleting antibody (α-CD8), or (I) an isotype control antibody (IgG) or an anti-IL-17A blocking antibody (α-IL-17A). **(J)** Quantification of the epidermal tongue length at 5 days after wounding wild type or *Il17a^-/-^* mice exposed to BHI-tween or control mice. (G-J) Data are representative of two independent experiments. Each dot represents the measured length of an individual epidermal tongue. * p < 0.05; ** p < 0.01; *** p < 0.001; **** p < 0.0001; ns, not significant (two-tailed unpaired Student’s t test, G-J).

GO term enrichment analysis further reinforced the type 17 signature (including functional enrichment for cytokine-cytokine receptor interaction, inflammatory response, nuclear receptor transcription pathway and cellular response to xenobiotic stimulus) (**Fig. 5D**) and revealed that terms associated with tissue repair and epithelial regeneration (including morphogenesis of an epithelium, gland morphogenesis and tissue morphogenesis) were enriched in detergent-induced Tc17 cells. Specifically, these terms were driven by genes such as *Ccr6*, *Fzd4*, *Areg*, *Furin*, and *Itgav* (**Fig. 5D**). Moreover, superimposing a previously published tissue repair gene set (*5*) onto both the bulk and single cell RNA-seq data revealed a strong enrichment of the tissue repair signature in CD8^+^ T cells from mice treated with mild detergent, whereas this signature was completely absent in HSV-1-induced CD8^+^ T cells (**Fig. 5E, F**).

Taken together, these data confirm that CD8^+^ T cells responding to detergent exposure are skewed towards a Tc17 phenotype, and their transcriptional signature supports the idea that these cells may have a reparative function.

### Wound repair is accelerated by xenobiotic-induced CD8^+^ T cells in an IL17A dependent manner

To test the potential wound repair property of detergent-elicited CD8^+^ T cells, we used an established method of back skin punch biopsy, which allows for the measurement of epithelial regeneration into the wound bed (*5–8*). Mice treated with mild detergent prior to back skin wounding exhibited significantly accelerated wound repair, as indicated by increased epithelial tongue length 5 days after wounding (**Fig. 5G**). Importantly, detergent-accelerated wound repair was dependent on CD8^+^ T cells (**Fig. 5H** and **fig. S5D**) and IL-17A (**Fig. 5I, J and fig. S5E, F**). Thus detergent-induced CD8^+^ T cell accumulation, and IL-17A production are sufficient to accelerate wound repair in the skin.

Collectively, this works suggests that detergent-exposed skin co-opts endogenous retroelements in Langerhans cells to preemptively elicit pro-repair tissue-resident immunity, an integral feature of barrier maintenance and host preservation.

## Discussion

Here, we propose that active recognition of retroelements by the immune system contributes to barrier tissue protection and regeneration.

While adaptive immunity to EREs has been described in the context of autoimmune disorders or cancer (*14, 15*), our present work uncovers that these responses can also occur in the context of benign settings and more specifically, in response to xenobiotics such as mild detergent. This observation raises the intriguing question about the tissue- and cell-specific control of these elements in both health and disease. The acquisition of EREs has required the integration of molecular mechanisms to prevent aberrant expression and protect the host from these genetic parasites (*40, 41*). Further, immune reactivity to EREs is under the control of negative selection and, as we further confirm here, pressure from regulatory T cells. However, in a similar manner to what has been previously described for tissue-specific antigens, some of these elements are not expressed by mTECs and as such, can escape negative selection (*42–44*). More specifically, and in agreement with previous reports, we uncover a remarkable partitioning of tissue and cell specific expression of EREs (*10, 45*). Of particular interest is the heightened expression of EREs within Langerhans cells. This dominant population of antigen presenting cells within the skin forms a dense network within the epidermis, uniquely positioning them as primary sentinels of the environment (*1, 46*). Our work proposes that ERE expression by Langerhans cells could act as a powerful interpreter of environmental stressors and help calibrate tissue resilience. This previously unappreciated role for Langerhans cell in mediating adaptive responses to retroelements, a phenomenon that promotes tissue regeneration, sheds an important light onto the physiological relevance of this dominant population of antigen presenting cells.

Our results also propose that adaptive immunity to retroelements is not exclusive to pathogenic states, and that reactivation caused by mild environmental exposure may positively contribute to the multilayer network of responses involved in tissue protection. The adaptive immune system is fundamentally malleable to the local environment and readily expandable via expression of defined antigen receptors, making this system a formidable and long-lived force to protect tissues. Thus, adaptive immunity to EREs represents an important class of immunity able to synergize with the previously described immunity to the microbiota (*5–9, 23, 24*), to promote tissue protection.

Over the past few decades, exposure to environmental stressors, such as cosmetics, detergents, and pollutants, at all barrier tissues has dramatically increased, along with a concomitant rise in chronic diseases, particularly in industrialized and westernized regions (*47–49*). Whether, under these contexts, aberrant reactivity to EREs and associated adaptive immunity contribute to inflammation and cancer remains to be investigated.

Our observation that adaptive immunity to EREs can be coopted by the host to protect barrier tissues supplies another layer to our understanding of the fascinating role of EREs in host physiology and evolution, and yet another lens through which to understand tissue physiology and disease.

## Acknowledgements

We thank K. Beacht, E. Lewis, G. Koroleva, Dr. P. Juliana Perez-Chaparro, the NIAID animal facility, and the NIAID Microbiome Program’s gnotobiotic animal facility for technical support. We thank Julie Laux and the NIAID Flow Cytometry Section for diligent assistance in all FACS sorting assays. Finally, we thank all Belkaid laboratory members for their generous support and providing feedback on this project, and particularly thank Dr. T. Farley and Dr. D. Corral for critical reading of this manuscript.

## Author Contributions

Conceptualization: YB, ACW

Methodology: YB, ACW, DSJ-L, VML, MS, SRK, LC, ES, CAR, AT, NB

Investigation: YB, ACW, DSJ-L, VML, MS, SRK, LC

Visualization: YB, ACW, DSJ-L, VML

Supervision: YB, ACW, VML

Writing—original draft: YB, ACW

Writing—review and editing: YB, ACW, DSJ-L, VML, SRK, LC, ES, CAR, AT, NB

## Data and materials availability

RNA-seq and scRNA-seq data were deposited into the Gene Expression Omnibus (GEO) data repository (GSE266527 for whole tissue bulk RNA-seq; GSE266528 for CD8^+^ T cell scRNA-seq; GSE266531 for cDC/LC/KC bulk RNA-seq; GSE266538 for CD8^+^ T cell bulk RNA-seq). All data are available in the main text or the supplementary materials.

## Materials and Methods

### Mice

C57BL/6NTac, C57BL/6J CD45.1 B6.SJL-*Ptprc^a^ Pepc^b^*/BoyJ (JAX: 002014), Ccr7^-/-^ B6.129P2(C)-*Ccr7^tm1Rfor^*/J (JAX: 006621), b2m^-/-^ C57BL/6NTac-[KO]beta2m, Il17a^-/-^ C57BL/6-[KO]IL17A were bred, maintained, and obtained by the NIAID Taconic Exchange Program. C57BL/6J (JAX: 000664) were obtained from The Jackson Laboratory. Germ free C57BL/6NTac mice were bred and maintained in the National Allergy and Infectious Diseases (NIAID) Microbiome Program gnotobiotic animal facility. Foxp3^YFPCre^Gata3^fl/fl^, huLangerin-DTR^EGFP^ B6.129S2-*Cd207^tm3.1(HBEGF/EGFP)Mal^*/J (JAX: 016940), H2-M3^-/-^ were bred and maintained at an American Association for the Accreditation of Laboratory Animal Care (AAALAC)-accredited facility at NIAID. Animals were housed in accordance with procedures outlined in the Guide for the Care and Use of Laboratory Animals. For experiments involving mouse strains bred and maintained at NIAID, littermate controls were used. For strains bred and maintained by the Taconic Exchange Program, wild type and knockout animals were cohoused at 3-5 weeks of age for 2-3 weeks prior to the start of experimental manipulations. Sex- and age-matched mice between 6-12 weeks of age were used for each experiment. All experiments were performed at NIAID under an Animal Study Proposal (LHIM-3E) approved by the NIAID Animal Care and Use Committee.

### Detergent preparation and exposure of mice

Sterile brain-heart infusion broth was emulsified with 1% Tween-80 (Sigma-Aldrich) or sodium dodecyl sulfate (Sigma-Aldrich) by vortexing. The emulsification was further sterilized by filtering through a 0.2 μm PES vacuum filter. For topical detergent exposure, each mouse was swabbed with ∼2mL of BHI-detergent emulsification across the surface of the ear pinnae or entire back skin using a sterile cotton swab. Application of detergent was repeated every other day for a total of 4 exposures, or performed daily for 7 days as indicated before analysis.

### Diet studies

The high fat diet (TD.06414; 60% of total calories from fat) and the corresponding control diet (TD.150064; 10% of total calories from fat) were both purchased from Envigo Teklad Diets. Immediately at weaning (3 weeks) mice were placed on the corresponding diet for 6 weeks before analysis.

### Murine tissue processing

Prior to tissue retrieval and processing, mice were euthanized with CO_2_.

#### Ear pinnae

Cells were isolated from ear pinnae as previously described (*23*). Briefly, ears were removed, separated into ventral and dorsal sheets, and digested in RPMI 1640 media (supplemented with 2mM L-glutamine, 1 mM sodium pyruvate and non-essential amino acids, 55 mM β-mercaptoethanol, 20 mM HEPES, 100 U/mL penicillin, 100 mg/mL streptomycin) with 0.5 mg/mL DNase I and 0.25 mg/mL Liberase TL purified enzyme blend at 37°C for 1 hour and 45 minutes. Digested ears were homogenized using sterile medicons (BD Biosciences) and subsequently filtered through a 70 μm cell strainer. To separate the epidermis from the dermis, were digested at 37°C for 45 minutes with 500 CU Dispase (Becton Dickinson) in HBSS without calcium and magnesium. The epidermis was then peeled from the dermis using curved forceps, then successively cut with scissors and further digested in RPMI 1640 media (supplemented as above) with 0.5 mg/mL DNase I and 0.05 mg/mL Liberase TL purified enzyme blend at 37°C for 1 hour and 10 minutes. Cells were then homogenized by pipetting up and down, and then passed through a 70 μm cell strainer.

#### Back skin

Back skin was cut away from shaved mice, and fat scraped off using a scalpel. 2 pieces of 1 cm^2^ back skin were digested in RPMI 1640 media (supplemented as above) with 0.5 mg/mL DNase I and 0.05 mg/mL Liberase TL purified enzyme blend at 37°C for 2 hours.

Digested skin was then homogenized using sterile medicons (BD Biosciences) and subsequently filtered through a 70 μm cell strainer.

Cervical lymph node: For downstream analysis of myeloid populations, lymph nodes were gently smashed using the plunger of a sterile 3mL syringe and digested in RMPI 1640 media (supplemented as above) with 0.5 mg/mL DNase I and 0.05 mg/mL Liberase TL purified enzyme blend at 37°C for 20 minutes. Digested lymph nodes were homogenized by gentle smashing with a plunger through a 70 μm cell strainer. For downstream analysis of lymphoid populations, lymph nodes were gently smashed with a plunger through a 70 μm cell strainer. Spleen: Spleens were first chopped into smaller pieces using scissors and then gently smashed through a 70 μm cell strainer with a plunger. Red blood cells in the resulting single cell suspensions were lysed using ACK lysis buffer for 5 minutes at room temperature.

### In vitro lymphocyte restimulation

To assess cytokine production potential, single cell suspensions from murine tissues were cultured *ex vivo* at 37°C for 2.5 hours in RPMI 1640 media (supplemented as above and with 10% fetal bovine serum) with 50 ng/mL phorbol myristate acetate (PMA) (Sigma-Aldrich), 5 μg/mL Ionomycin (Sigma-Aldrich), and 1:1000 dilution of GolgiPlug (BD Biosciences).

### Flow cytometry analysis

Single cell suspensions were incubated with fluorophore-conjugated antibodies against surface markers for 30 minutes at 4°C in PBS, unless otherwise noted. LIVE/DEAD Fixable Blue Dead Cell Stain Kit (Invitrogen) was used to exclude dead cells. Cells were fixed and permeabilized with the FoxP3/ Transcription Factor Staining Buffer Set (eBioscience) according to the manufacturer’s instructions. Cells were then stained with fluorophore-conjugated antibodies against intracellular markers for at least 30 minutes at 4°C, unless otherwise noted. All staining was performed in the presence of purified anti-mouse CD16/32 (clone 93). All antibodies used are listed in table S2.

For detection of LINE-1 ORF1p, fixed and permeabilized cells were stained with the anti-ORF1p antibody for 1 hour at room temperature. For detection of the MLV envelope, the hybridoma which generates the 83A25 monoclonal antibody that recognizes ecotropic and non-ecotropic ERV envelope protein (MLV Xmv45) was acquired from Leonard Evans (*50*). Single cell suspensions were incubated with hybridoma 83A25 supernatant for 1 hour at room temperature in HBSS, followed by staining with a biotinylated anti-rat IgG2A antibody (clone RG7/1.30, BD Biosciences), and finally streptavidin-PE-Cy7 (BioLegend).For LINE-1 ORF1p and MLV envelope, integrated mean fluorescence intensity was calculated by multiplying the frequency of positive cells by the MFI of positive cells, as determined by an isotype control.

### Ear thickness measurement

A digital caliper (Mitutoyo) was used to measure ear thickness. The change in ear thickness over time was reported as relative to the ear thickness of mice before detergent exposure.

### Parabiosis

Parabiosis was performed as previously described (*51*). Briefly, congenically-mismatched, weight-matched mice were anesthetized with ketamine/ xylazine (10 μg/g body weight) and shaved on the right or left side body as needed. Incisions were created from the olecranon to the knee joint. Parabiotic pairs were then created by suturing the olecranon and knee joints. Dorsal and ventral skin was then attached by continuous stapling and suturing as needed. Parabiotic pairs were left conjoined for 30 days before analysis.

### Confocal microscopy

Mouse ears were cut and split into ventral and dorsal sheets, with any remaining cartilage removed via scraping under a light dissecting scope. Ears were then fixed overnight at 4°C in 1% paraformaldehyde (in PBS), washed, and blocked in a buffer of 1% bovine serum albumin and 0.25% Triton X-100 overnight at 4°C. Antibodies in blocking buffer were added to samples to stain for 24-36h at 4°C. Samples were washed in PBS, mounted in ProLong Gold, and dried at room temperature overnight prior to acquisition. Images were captured with a Leica TCS SP8 confocal microscope fitted with HyD and PMT detectors using a 40X oil objective (HC PL APO 40X/1.3 oil). Images were analyzed using Imaris software (Bitplane).

### Tetramer staining and enrichment

Staining for all tetramers used (MAIT, MR-1:5-OP-RU PE; YopE, MHC-I:SVIGFIQRM PE and APC; LINE-1, MHC-I:KSINVIHYI PE and APC, and MHC-1:ISLMNINAKIL PE and APC) were performed at room temperature for 1 hour. Enrichment of tetramer+ cells was performed as previously described (*5*). Briefly, using anti-PE or anti-APC coupled magnetic beads, samples were enriched for bead-bound cells on magnetized columns (StemCell Technologies #17684 and #17681, respectively). Samples then underwent surface staining on ice with antibodies as described above. To calculate frequencies of enriched tetramer^+^ CD8^+^ T cells, the frequency of tetramer^+^ cells was divided by the frequency of enrichment (determined by the pre- and post-enrichment tetramer^+^ cell frequency).

### ERE peptide prediction and pooling

ERE loci with a minimum expression level of 4 TPM and a log_2_ Fold Change of > 2 with a false discovery rate of < 0.05 between BHI-tween-exposed and naïve Langerhans cells were identified by bulk RNA-seq as described above. Loci which were expressed over 4 TPM in medullary thymic epithelial cells (Chr5 2.4- and Chr17 3.2+) were excluded from peptide prediction analysis to limit the potential identification of highly autoreactive peptides. The amino acid sequences of the remaining ERE loci were attained using the Genome-based Endogenous Viral Element Database (gEVE version 1.1, Nakagawa 2016) Mmus38 open reading frame amino acid sequences. The amino acid sequences were analyzed for MHC-I binding epitopes using the Immune Epitope Database & Tools MHC-I binding predictions version 2.24 with the netMHCPan 4.1 EL prediction method for both H2-D^b^ and H2-K^b^. Predicted peptides with a score of > 0.2 and a rank of < 0.5 were considered positive MHC-I binding epitopes. Peptides were ordered from Genscript at crude purity for pool stimulation, as described below. Peptides were pooled into groups of 12 initially. Then, through subsequent rounds of stimulation (with freshly isolated CD8 T cells and purified DCs each time), pools which yielded positive cytokine production were broken down into smaller pools until eventually individual peptides were identified. Once individual peptides were identified, they were ordered at ≥ 98% purity for stimulation.

### Ex vivo ERE peptide stimulation/ DC-T cell co-culture assay

CD45+ CD90.2+ TCRβ+ CD8β+ T cells were sort-purified on a FACSAria from ear pinnae skin of mice exposed to BHI-tween 14 days prior. For DC purification, single cell suspensions from lymph nodes and spleens were magnetically enriched for CD11c+ cells using CD11c microbeads and MACS separation columns according to the manufacturer’s recommendations (Miltenyi Biotec). Purified splenic DCs were then pulsed with peptides at a final concentration of 15 μg/mL/peptide for 30 minutes at 37°C with 5% CO_2_. Peptide pulsed DCs and sort purified CD8β+ T cells were co-cultured at a 15:1 ratio (2-5X10^3^ CD8β+ T cells) in a 96 well U-bottom plate in 10% RPMI (supplemented as described above) for 16 hours at 37°C with 5% CO_2_.

GolgiPlug was added for the final 4 hours of culture, after which cells were assessed for cytokine potential by intracellular flow cytometry as described above.

### FACS sort purification of lymphocytes and keratinocytes

To sort CD8+ T cells, single cell suspensions from ear pinnae skin of WT mice exposed to BHI-tween 14 days prior, or infected with HSV-1 7 days prior were stained with antibodies against CD45, lineage (CD11b, NK1.1, γδTCR), CD90.2, TCRβ, CD4, CD8β, and DAPI for 30 minutes at 4°C. CD8+ T cells were sorted as DAPI-lineage-CD45+ CD90.2+ TCRβ**+** CD4-CD8β on a FACSAria. For *ex vivo* peptide restimulation assays, cells were sorted into 10% RPMI (supplemented as above). For bulk RNA-sequencing, cells were sorted directly into lysis buffer and immediately placed on dry ice, then stored at −80°C. For single cell RNA-sequencing, cells were additionally stained with TotalSeqA hashtag antibodies (BioLegend), sorted into 10% RPMI (supplemented as above), then washed and counted before pooling and loading into the 10X Chromium Controller, as described below. To sort keratinocytes, dendritic cells, and Langerhans cells, single cell suspensions from ear pinnae skin of control mice or mice exposed to BHI-tween daily for 7 days were stained with antibodies against Sca-1, CD64, CD31, CD34, CD49f, CD11c, CD45, CD24, CD11b, Ly6G, lineage (NK1.1, TCRβ, γδTCR) and DAPI for 30 minutes at 4°C. Keratinocytes were sorted as DAPI-lineage-Ly6G-CD45-CD34-CD49f+ Sca-1+. Conventional DC1 were sorted as DAPI-lineage-Ly6G-CD11c+ CD64-CD11b-CD24+, and conventional DC2 were sorted as DAPI-lineage-Ly6G-CD11c+ CD64-CD11b+ CD24-, then pooled during collection. Langerhans cells were sorted as DAPI-lineage-Ly6G-CD11c+ CD64-CD11b+ CD24+. Sorting was performed on a FACSAria directly into lysis buffer, immediately placed on dry ice, then stored at −80°C.

### Diphtheria toxin treatment

For full diphtheria toxin (DTX) treatment, huLangerin-DTR mice were injected intraperitoneally (i.p.) with 200 μg of DTX 3 days before the start of BHI-tween exposure. Then, 10 days after the first DTX treatment, mice were injected i.p. with 100 ng of DTX every 4 days.

### Single cell RNA-seq

For single cell RNA-seq, CD8+ T-cells from the mouse ear were sorted with a FACSAria sorter from samples treated with BHI-tween or HSV-1 and labeled with TotalSeqC hashtags antibodies (Biolegend). All samples were pooled together, and 40,000 cells were loaded to a Chromium Single Cell Controller (10X Genomics) to encapsulate cells into droplets. A total of 4 10X lanes were run, resulting in a total of 40,000 loaded cells. Libraries were prepared using a Chromium Single Cell 5’ Reagent kits following manufacturer’s instructions. Libraries were then sequenced on an Illumina NextSeq 2000 (NextSeq 2000 P3 Reagents (200 Cycles), Illumina). Illumina files were converted to FASTQ files using bcl-output command and data were filtered and mapped to mm10 reference genome by using cellranger 7.0.0 (10X Genomics).

The overall sequencing quality was high in the four T cell mRNA libraries with > 93% of bases in the barcode and UMI regions had Q30 quality score or above, and > 90% of bases in the RNA reads had Q30 or above. Furthermore, the median gene count range was 2,137 – 2,471 along with a sequencing saturation of > 50% and about 70% of the reads mapped confidently to the transcriptome. The mean read count per cell was ∼15,000 across the four samples. For the four TCR libraries, the percentage of cells with productive V-J spanning (TRA, TRB) pairs was > 80% and the mean read count per cell was between 4,000 – 6,000. For the four HTO libraries, mean read count per cells was between 1,900 – 2,300 with > 97% of valid barcodes.

Data were normalized in Seurat (v. 5.0.2) (*53*). Cells with 200 – 4000 RNAs, less than 5% of mitochondrial RNA and less than 20,000 RNA reads were kept for further analysis. Hashtag libraries were demultiplexed using Seurat’s HTODemux and negative and doublet cells were filtered out. In the first round of data processing 14 PCs were utilized for neighbor finding and a resolution of 0.8 for cluster finding. Following this step cell types were automatically assigned using scType (*54*) and cells assigned as “Cancer cells”, “Endothelial”, “Erythroid-like and erythroid precursor cells”, “Myeloid Dendritic cells” were filtered out. Data was re-processed using 13 PCs for neighbor finding and a resolution of 0.8 for cluster finding and cell types were assigned manually. TCR data was processed using scRepertoire (v. 1.10.1) using the aa clonotype definition and added to the Seurat object. To visualize Tc17, Tc1 and tissue repair signatures on the UMAP plot a score was calculated based on a subset of genes using UCell (v. 2.4.0) (*54*).

### Bulk RNA-seq on sorted cell populations

For bulk RNA-seq on sorted cell populations we sorted CD8+ T cells from the ear of mice treated with HSV-1 or BHI-tween, as well as Langerhans Cells, cDC1, cDC2, and keratinocytes from the ear of unassociated mice and mice treated with BHI-tween using a FACS Aria sorter. cDC1 and cDC2 samples were combined post-sort for RNA extraction. Then, RNA was extracted using the Qiagen miRNeasy kit, and sequencing library was prepared using Tecan SoLo total RNA kit for mouse. Libraries were sequenced as 1 x 150bp reads on a Illumina NextSeq 2000 using the P3 200 cycle kit.

### Whole tissue RNA-seq

Whole tissue ear RNA-seq was performed utilizing a 4mm punch biopsy of mice ears. RNA was extracted utilizing the Qiagen RNeasy kit for tissue. Sequencing libraries were generated using the Illumina Stranded Total RNA Prep, Ligation with Ribo-Zero Plus kit, according to the manufacturer’s instruction. Final libraries were sequenced as paired end 2 X 150 bp reads on the NovaSeq 6000 instrument using the SP 300 cycle sequencing kit.

### Bulk RNA-seq analysis

Sequencing reads were mapped to the C57BL/6 mouse genome (GRCm38: mm10) and differential gene expression was calculated utilizing homer’s getDifferentialExpression with default parameters (*55*). For ERV analysis, homer’s analyzeRepeats with a custom ERV file was utilized, as well as with the parameter repeats. Differentially expressed genes (FDR < 0.05, log2FC < 2) were used for gene ontology (GO) enrichment analysis with MetaScape (*56*).

### Bulk RNA-seq endogenous retroelement (ERE) analysis

Endogenous retroelement expression was determined from bulk RNA-sequencing as previously described (*8*). Briefly, sequencing reads were mapped to the C57BL/6 mouse genome (GRCm38: mm10) genome with STAR using stringent mapping parameters (-outFilterMultimapNmax2).

Retroelement expression was assessed using HOMER4.11 according to the annotation provided by gEVE version 1.1 (*52*). Reads for elements of ERE classes were analyzed using the analyzeRepeats function with parameters repeats mm10-d. Reads for ERE loci were also analyzed using the analyzeRepeats function, but with parameters mm10-count genes-noadj-d. Differential expression was calculated with DESeq2. EREs with a False Discovery Rate of < 0.05 were considered differentially expressed. For whole tissue retroelement analysis, a log2 Fold Change > 1 was considered differentially expressed; for all other retroelement analysis, a log2 Fold Change > 2 was considered differentially expressed.

### Back skin wounding and epifluorescence microscopy of wound tissue

Wounding and quantification of wound healing were performed as previously described (*5–9*). Male mice in the telogen phase of hair growth (10-12 weeks old) were exposed to BHI-tween. 12 days after the initial exposure, exposed and control animals were anesthetized with ketamine/xylazine, and back skin was shaved with clippers and depilated with Nair. A 6mm punch biopsy tool was used to perforate skin, and iris scissors used to remove the epidermis and dermis, creating a circular full thickness wound. 5 days after wounding, back skin tissue was removed, fixed in 4% PFA for 4 hours at 4°C, washed in PBS twice, and incubated overnight in 30% sucrose at 4°C. Fixed wounds were embedded in OCT compound (TissueTek), frozen on dry ice and cryosectioned into 20 μm slices on SuperFrost Plus microscope slides (Fisher). Extra OCT was scraped off of dried slides, which were then fixed in 4% PFA for 10 minutes at room temperature, washed with PBS, permeabilized with 0.1% Triton X-100 for 10 minutes at room temperature, washed with PBS, and blocked for 1 hour in blocking buffer (2.5% Normal Goat Serum, 1% BSA, 0.3% Triton X-100 in PBS) at room temperature. Sections were then incubated overnight at 4°C with the primary antibody chicken anti-mouse Keratin 14 (Poly9060, BioLegend) at a 1:400 dilution in blocking buffer with anti-CD16/32. After washing with PBS, sections were incubated with the secondary antibody goat anti-chicken conjugated with AlexaFluor647 (Jackson ImmunoResearch) for 1-2 hours at room temperature. Slides were washed with PBS, stained with DAPI (1:100) 10 minutes at room temperature, washed again and mounted with ProLong Gold. Wound images were captured with a Leica DMI 6000 epifluorescence microscope fitted with a Leica DFC360X monochrome camera. Tiled and stitched images of wounds were collected with a 20X/0.4NA dry objective. Images were analyzed using Imaris software (Bitplane).

### In vivo treatment with blocking antibodies

For depletion of CD8 T cells, mice were injected intraperitoneally (i.p.) with 200 μg of anti-CD8 (2.43, BioXcell) or mouse IgG2b isotype control (LTF-2, BioXcell) daily for 3 days prior to the initial BHI-tween exposure. Following the initial BHI-tween exposure, mice were injected i.p. with with 200 μg of anti-CD8 or IgG1 every four days until the endpoint. For blocking of IL-17A, mice were injected i.p. with 0.5 mg of anti-IL-17A (17F3, BioXcell) or mouse IgG1 isotype control (MOPC-21, BioXcell) two days prior to the first BHI-tween exposure, and then every other day until the endpoint.

### HSV-1 infection

Mice were infected in the dorsal ear pinnae as previously described (*57*). Briefly, mice were anesthetized with isoflurane, then 1X10^6^ PFU of HSV-1 was pipetted onto the dorsal ear pinnae, which was subsequently poked 5 times with a bifurcated needle (Roechling Medical).

### Statistical analysis

Groups were compared with PRISM software (Versiona 9.5.1). Stastical methods used are specified in the figure legends.

---

**Fig. S1.**
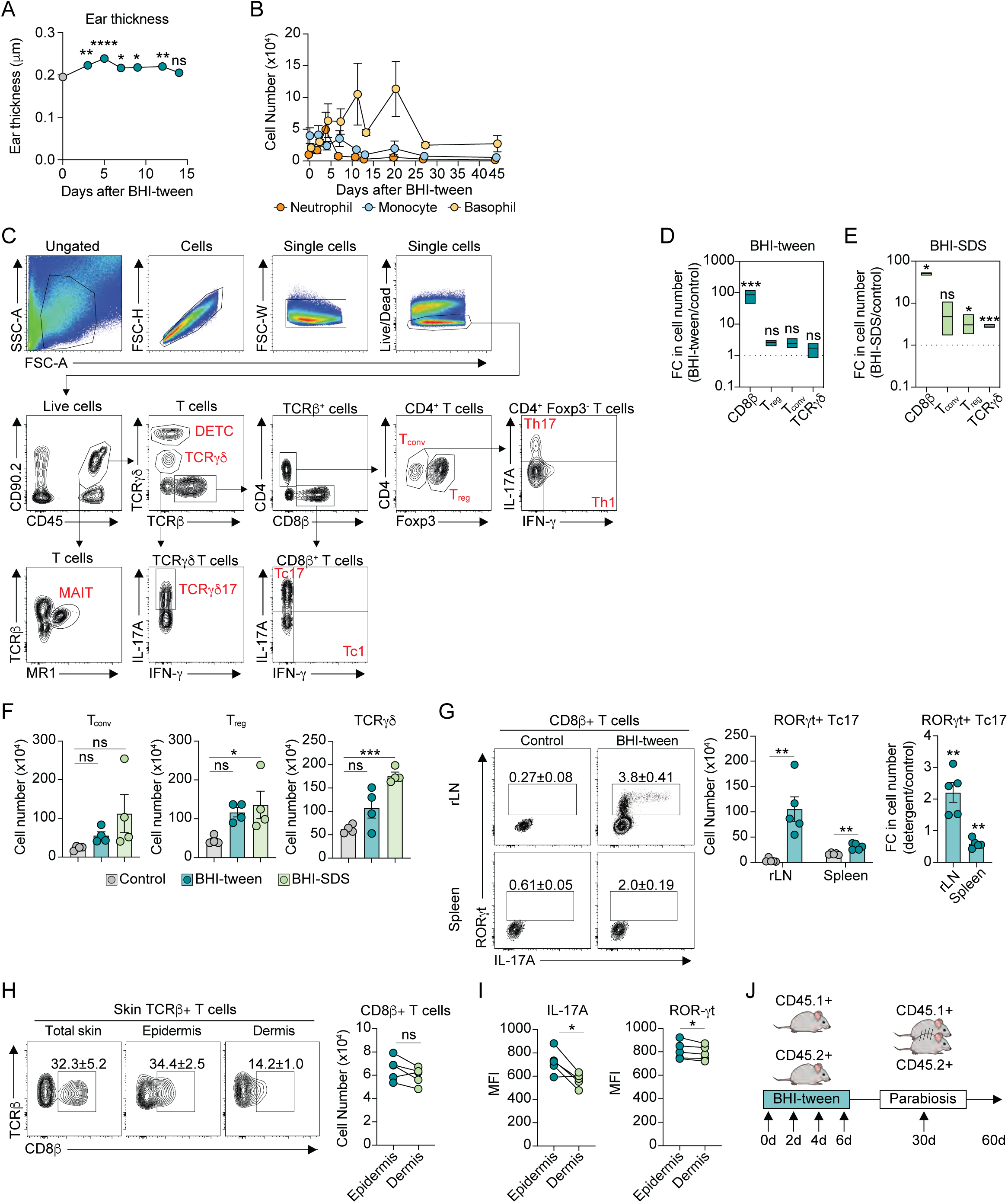
Immune responses to disparate environmental exposures are unique and occur in the absence of inflammation. **(A)** Ear thickness measured in microns (μm) in control or BHI-tween exposed mice at the indicated time. **(B)** Fold change in absolute numbers of neutrophils, monocytes, and basophils in ear pinnae skin at the indicated time after BHI-tween exposure. **(C)** Gating strategy to identify various lymphocyte populations in the skin. DETC, dendritic epidermal T cell; T_conv_, conventional T cell (CD4^+^ Foxp3^-^); T_reg_, regulatory T cell (CD4^+^ Foxp3^+^); MAIT, mucosal associated invariant T cell. **(D-E)** Fold change in absolute numbers of CD8β^+^, T_regs_ (CD4^+^ Foxp3^+^), T_conv_ (CD4^+^ Foxp3^-^), and TCRγδ-mid, in ear pinnae skin of (D) BHI-tween exposed mice, or (E) BHI-SDS exposed mice. **(F)** Absolute numbers of lymphocyte populations in BHI-tween or BHI-SDS exposed mice. **(G)** (Left) representative FACS plots showing frequencies or (right) absolute numbers of Tc17 cells (RORγt^+^ IL-17A^+^ CD8β^+^ T cells) in the ear pinnae-regional lymph node (rLN) or spleen of control or BHI-tween-exposed mice. **(H)** (Left) representative FACS plots showing frequencies or (right) absolute numbers of CD8β^+^ T cells in whole ear pinnae skin, ear pinnae epidermis, or ear pinnae dermis. **(I)** Mean fluorescence intensity of (left) IL-17A or (right) RORγt in CD8β^+^ T cells in ear pinnae epidermis or dermis. **(J)** Experimental design of parabiosis to assess tissue residency in response to detergent exposure. (A-H) Data are representative of at least two independent experiments. (A-D) Each dot or bar is the average of 5 individual mice. (E-H) Each dot represents an individual mouse. * p < 0.05; ** p < 0.01; *** p < 0.001; **** p < 0.0001; ns, not significant (one-way ANOVA with Dunnett’s multiple comparisons test (compared to control or day 0) for A and E; two-tailed unpaired Student’s t test for C, D, F; two-tailed paired Student’s t test for G-H).

**Fig. S2.**
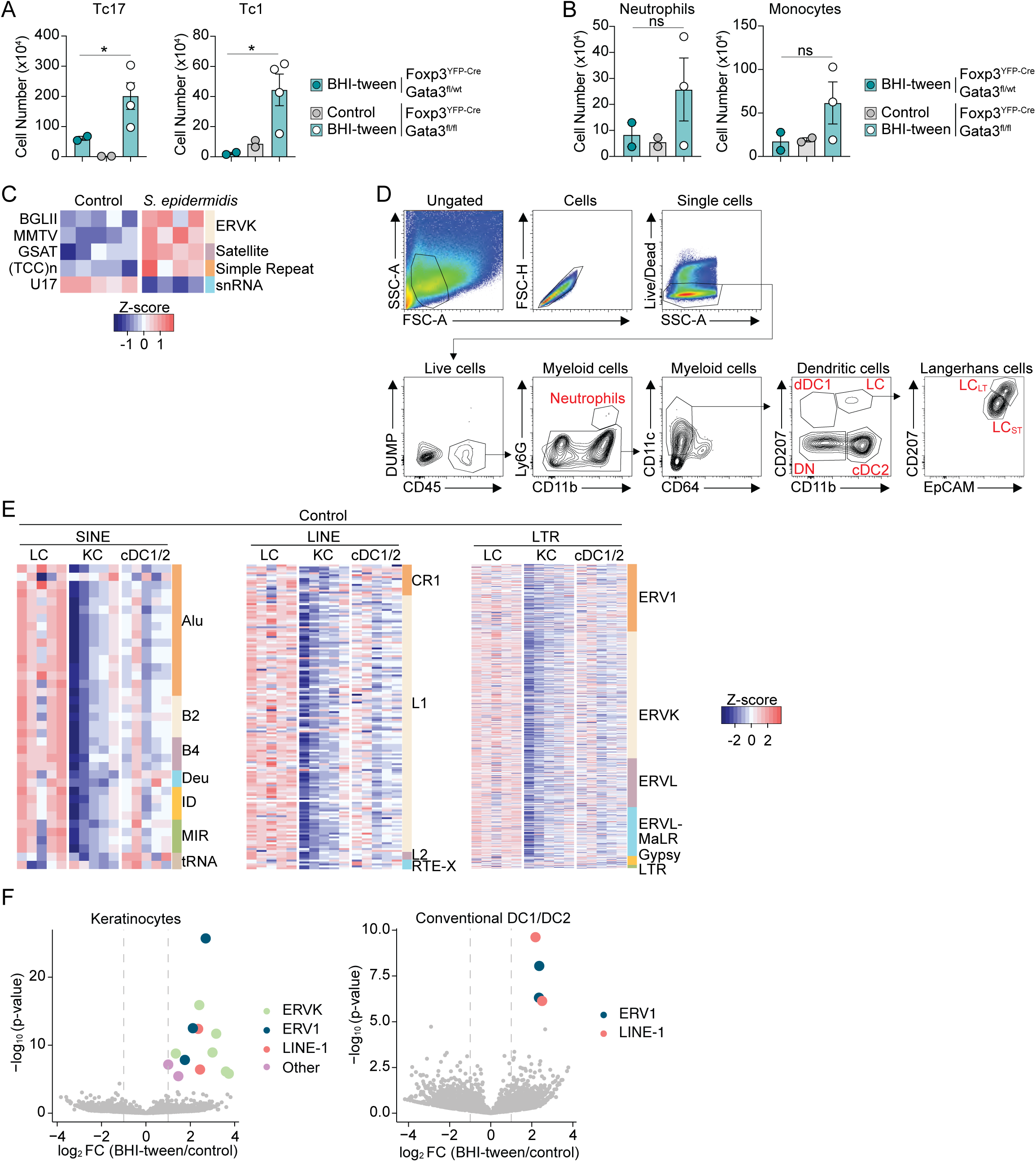
Endogenous retroelements are expressed in a context and cell-type specific manner. **(A)** Absolute numbers of (left) Tc17 cells, (right) Tc1 cells, **(B)** (left) neutrophils, or (right) monocytes in control or BHI-tween exposed Foxp3^YFP-Cre^Gata3^fl/wt^ (wild type) or Foxp3^YFP-Cre^Gata3^fl/fl^ mice. **(C)** Heatmap displaying fold change of significantly differentially expressed retroelement elements (FC > 2, FDR < 0.05) by RNA-sequencing of whole homogenized ear pinnae skin from control mice or animals exposed to *S. epidermidis* daily for 7 days. Retroelement elements are listed on the left, and retroelement classes on the right. **(D)** Gating strategy to identify dendritic cells and Langerhans cell subpopulations in the skin. dDC1, dermal type 1 dendritic cells; LC, Langerhans cells; cDC2, conventional type 2 dendritic cells; DN, double negative; LC_LT_, long-term LCs; LC_ST_, short-term LCs. **(E)** Heatmap displaying row z-score of retroelement elements from RNA-sequencing of naïve Langerhans cells (LC), keratinocytes (KC; Sca-1^+^, CD49f^+^), and conventional dendritic cells (cDC1/2) sort-purified from ear pinnae skin of control animals. (Left) Short-Interspersed Nuclear Elements (SINE), (middle) Long-Interspersed Nuclear Elements (LINE), (right) Long Terminal Repeats (LTR). Retroelement families are listed on the right. Deu, deuterostome; ID, identifier; MIR, mammalian-wide interspersed repeat; RTE, retrotransposable element clade; ERV, endogenous retrovirus; MaLR, mammalian apparent LTR retrotransposon. **(F)** Volcano plot displaying differentially expressed retroelement loci from RNA-sequencing of (left) bulk keratinocytes (Sca-1^+^, CD49f^+^), and (right) conventional dendritic cells sort-purified from ear pinnae skin of mice exposed daily to BHI-tween for 7 days as compared to control mice.

**Fig. S3.**
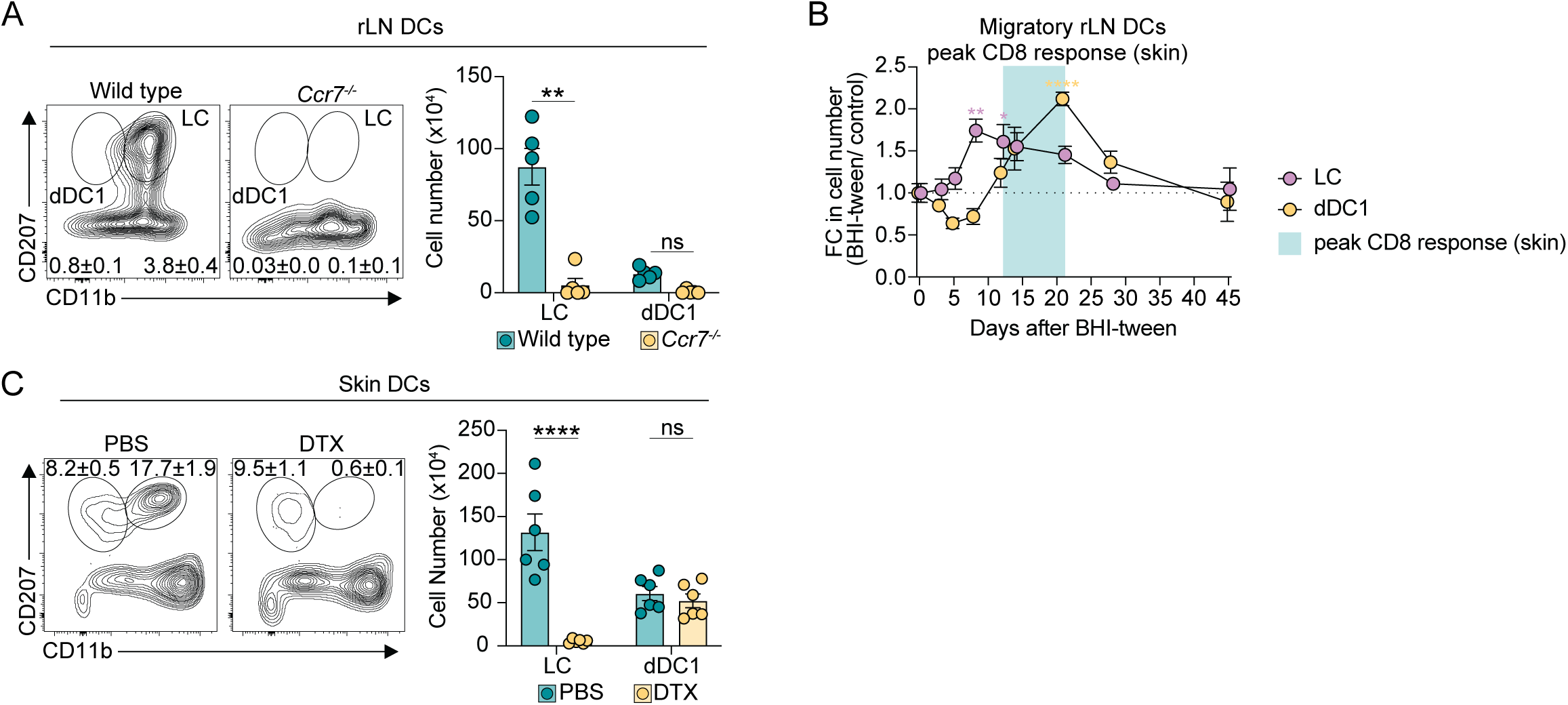
Langerhans cell migration to regional lymph nodes is required for the CD8^+^ T cell response to detergent. **(A)** (Left) representative FACS plots showing frequencies or (right) absolute numbers of Langerhans cells and dDC1s in ear pinnae-regional lymph nodes of wild type or *Ccr7^-/-^* animals treated either as controls or 14 days after BHI-tween exposure. **(B)** Fold change in absolute numbers of migratory dDC1s or Langerhans cells in the ear pinnae-regional lymph node of mice at the indicated time after BHI-tween exposure as compared to control mice. Peak accumulation of CD8β^+^ T cells in the skin is highlighted. **(C)** (Left) representative FACS plots showing frequencies or (right) absolute numbers of Langerhans cells and dDC1s in ear pinnae skin of BHI-tween exposed mice (day 14) treated with either PBS or diphtheria toxin (DTX). (A-C) Data are representative of at least two independent experiments. (A, C) Each dot represents an individual mouse. (B) Each dot is the average of 5 mice. * p < 0.05; ** p < 0.01; *** p < 0.001; **** p < 0.0001; ns, not significant (two-tailed unpaired Student’s t test for A, C; two-way ANOVA with Tukey’s multiple comparisons test for B).

**Fig. S4.**
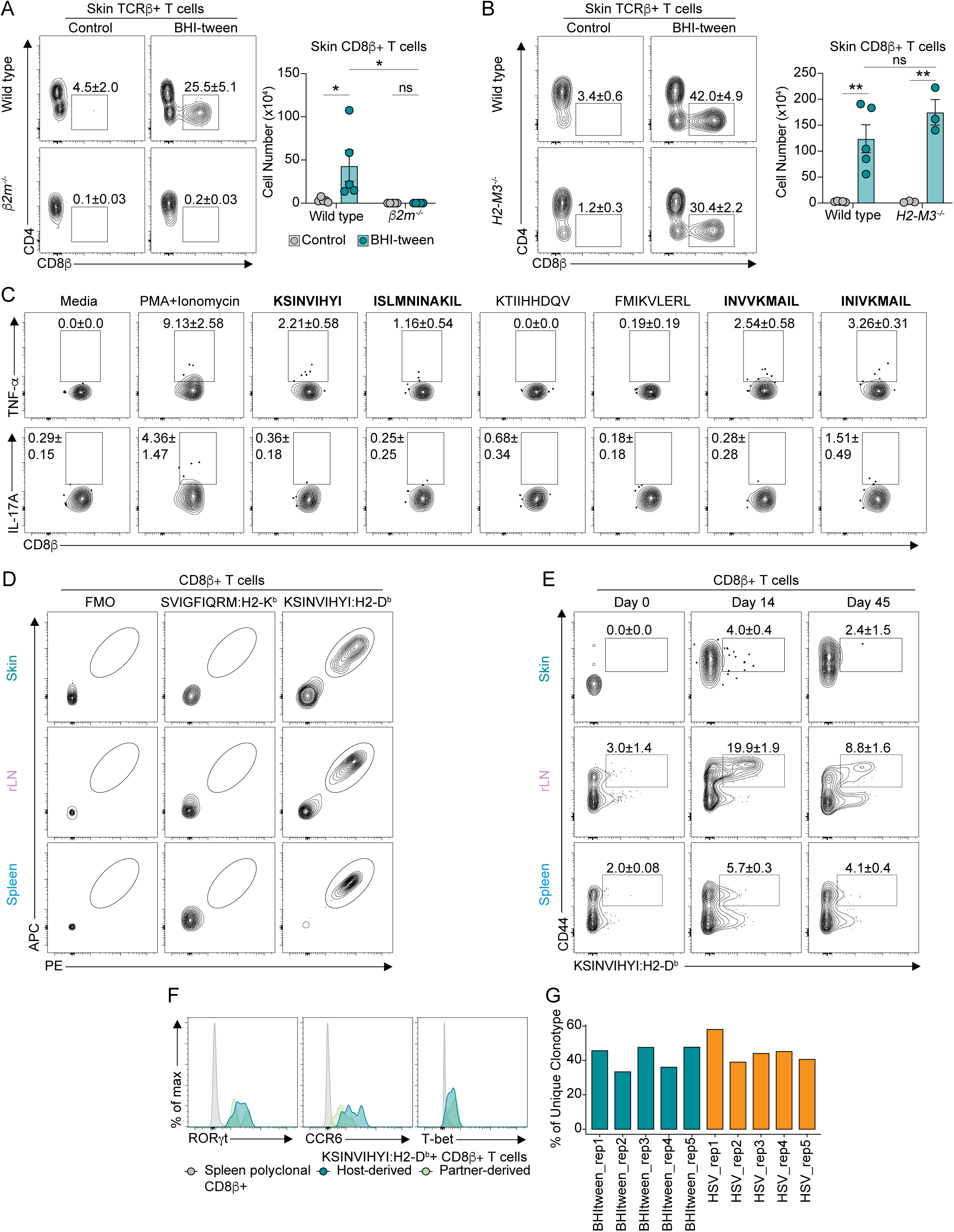
CD8^+^ T cells responding to detergent exposure recognize LINE-1-derived antigens. **(A)** (Left) representative FACS plots showing frequencies or (right) absolute numbers of CD8β^+^ T cells in ear pinnae skin of wild type or *β2m^-/-^* animals treated either as controls or 14 days after BHI-tween exposure. **(B)** (Left) representative FACS plots showing frequencies or (right) absolute numbers of CD8β^+^ T cells in ear pinnae skin of wild type or *H2-M3^-/-^* animals treated either as controls or with BHI-tween 14 days prior. **(C)** Representative FACS plots showing frequencies of (top) TNF-α or (bottom) IL-17A production by CD8β^+^ T cells sorted from ear pinnae skin exposed to BHI-tween 14 days prior and stimulated *in vitro* with the indicated peptide, or PMA+Ionomycin. **(D)** Representative FACS plots showing frequencies of CD8β^+^ T cells labeled with both a PE and an APC conjugated form of the indicated tetramer in (top) ear pinnae skin, (middle) ear pinnae-regional lymph node, or (bottom) spleen of mice exposed to BHI-tween 14 days prior. FMO, fluorescence minus one. **(E)** Representative FACS plots showing frequencies of CD44^+^ KSINVIHYI:H2-D^b+^ CD8β^+^ T cells in the (top) ear pinnae skin, (middle) ear pinnae-regional lymph node (bottom) or spleen of mice at the indicated time after BHI-tween exposure. **(F)** Representative histograms showing expression of RORγt, CCR6, and T-bet by either host-derived or partner-derived KSINVINHYI:H2-D^b+^ CD8β^+^ T cells in ear pinnae skin, or total splenic CD8β^+^ T cells of parabiotic mice. **(G)** Bar plot showing frequency of unique clonotypes in each replicate mouse. (A-F) Data are representative of at least two independent experiments. (A, B) Each dot represents an individual mouse. * p < 0.05; ** p < 0.01; ns, not significant (two-way ANOVA with Tukey’s multiple comparisons test for A, B).

**Fig. S5.**
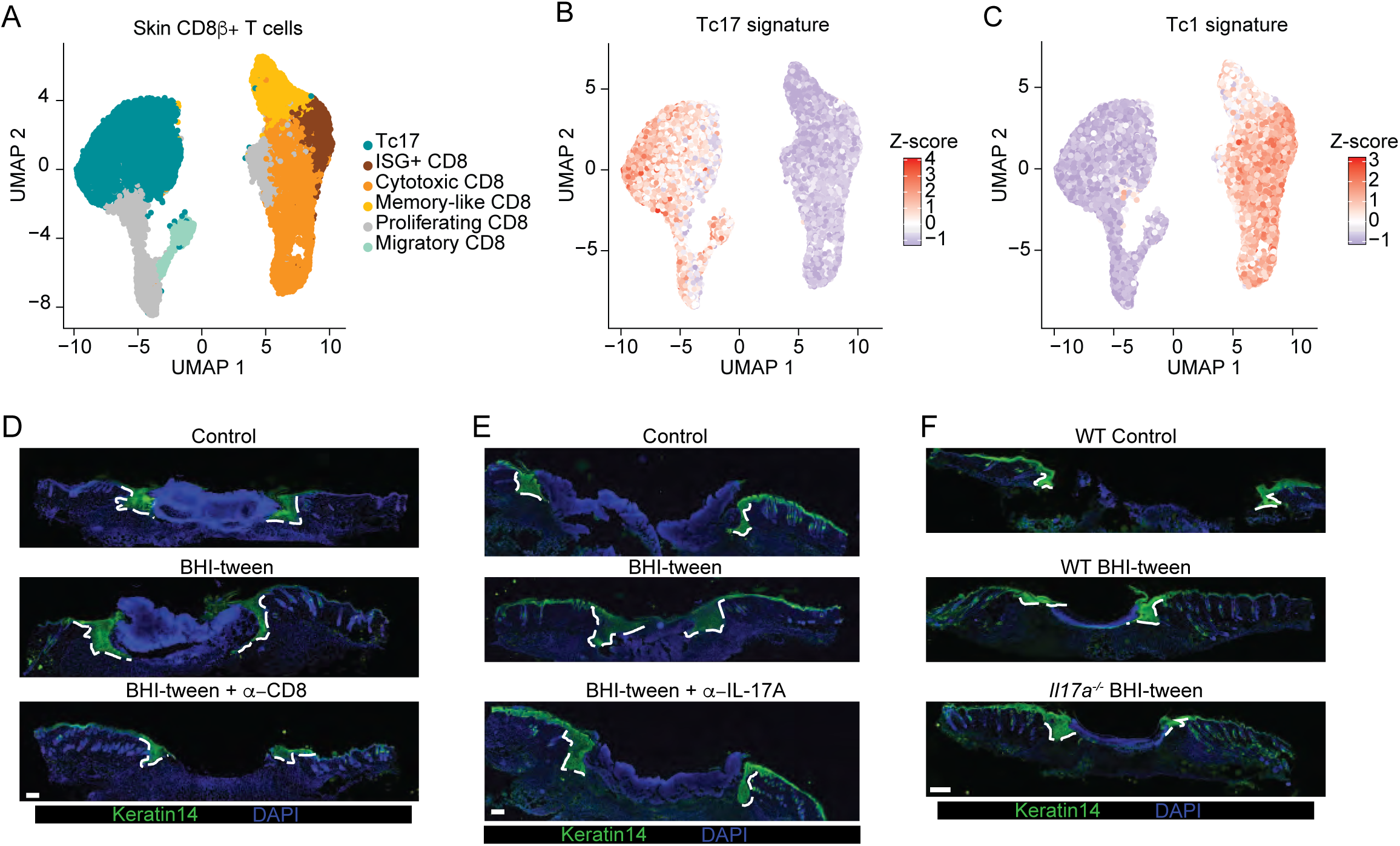
CD8^+^ T cells responding to detergent are transcriptionally heterogeneous and aid in wound repair. **(A-C)** CD8β^+^ T cells were sort-purified from ear pinnae skin of HSV-1-infected or BHI-tween-exposed animals for single cell RNA-sequencing analysis. (A) UMAP projection plot showing Seurat clusters, (B) the type 17 signature, or (C) the type 1 signature in ear pinnae skin CD8β^+^ T cells. **(D-F)** Mice were untreated or exposed to BHI-tween 12 days prior to back skin punch biopsy, and epithelial tongue length measured 5 days after wounding. (E) Representative immunofluorescence images of re-epithelialization in control mice, mice exposed to BHI-tween 12 days prior and treated with either IgG isotype control or anti-CD8α antibodies. Scale bars are 300 μm. (F) Representative immunofluorescence images of re-epithelialization in mice exposed to BHI-tween 12 days prior and treated with either IgG isotype control or anti-IL-17A antibodies. Scale bars are 200 μm. (G) Representative immunofluorescence images of re-epithelialization in wild type or *Il17a^-/-^* animals treated either as a control or with BHI-tween 12 days prior. Scale bars are 300 μm. (E-G) Data are representative of at two independent experiments.

**Table S1.**
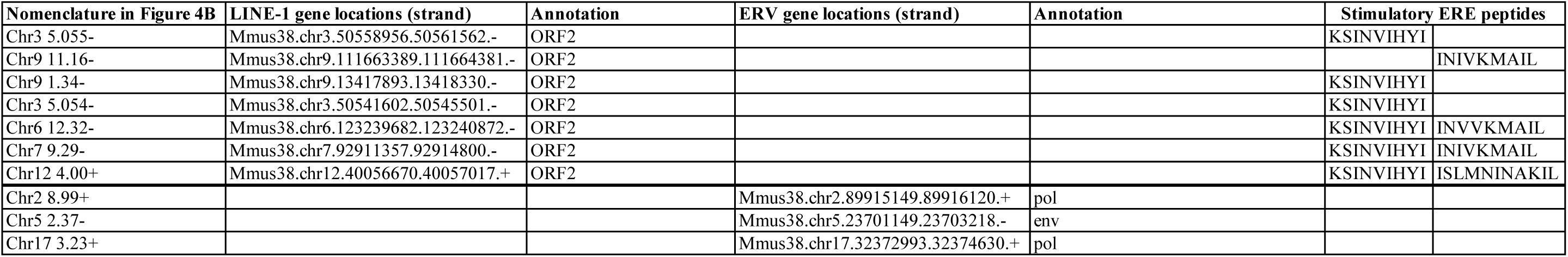
Chromosomal location and classification of endogenous retroelements differentially expressed in Langerhans cells after detergent exposure. Retroelement nomenclature is based on the chromosomal location and strand designation in GRCm38 as defined by gEVE database.

**Table S2.**
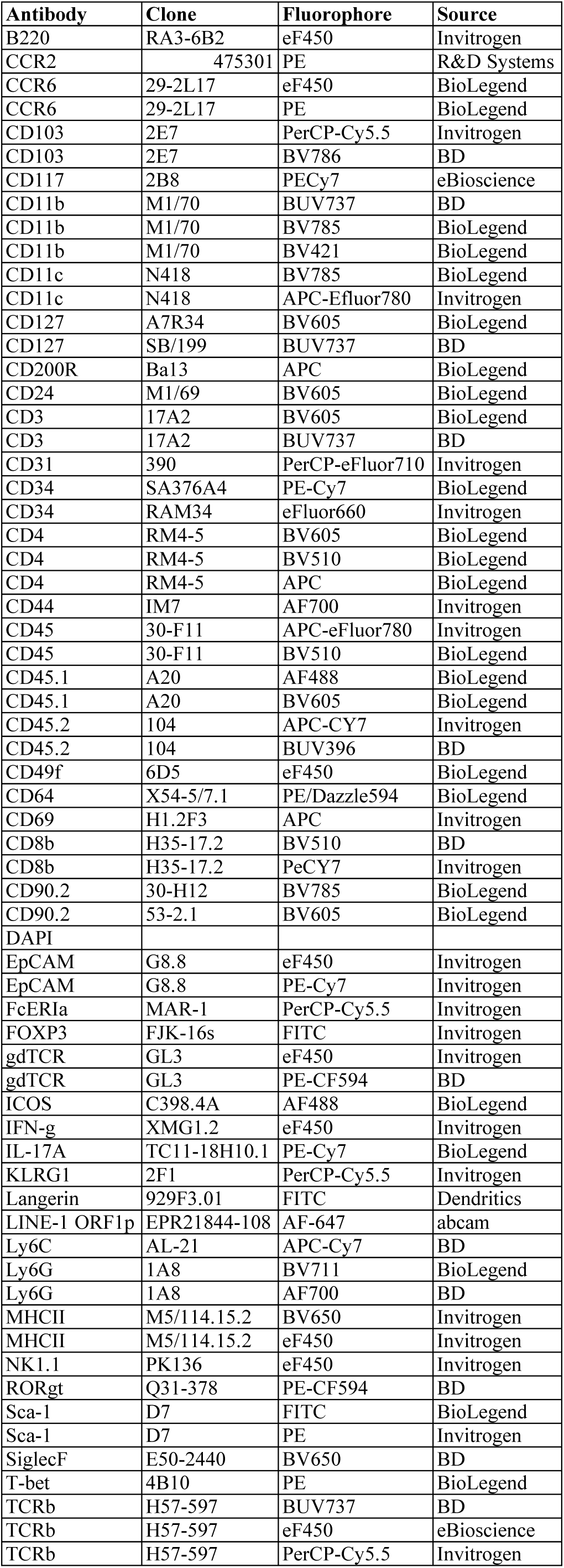
Antibodies used for flow cytometry. List of antibody specificity, clone, fluorophore and source for antibodies used in flow cytometry experiments.

